# Landscapes of ribosome dwell times and relationship with aminoacyl-tRNA levels in mammals

**DOI:** 10.1101/551838

**Authors:** Cédric Gobet, Benjamin Weger, Julien Marquis, Eva Martin, Frédéric Gachon, Felix Naef

## Abstract

Protein translation depends on mRNA-specific initiation, elongation and termination rates. While ribosome elongation is well studied in bacteria and yeast, less is known in higher eukaryotes. Here, we combined ribosome and tRNA profiling to investigate the relations between ribosome elongation rates, (aminoacyl-) tRNA levels, and codon usage in mammals. We modeled codon-specific ribosome dwell times and translation efficiencies from ribosome profiling, considering codon-pair interactions between ribosome sites. In mouse liver, the model revealed site and codon specific dwell times, as well as codon-pair interactions clustering by amino acids. While translation efficiencies varied significantly across diurnal time and feeding regimen, codon dwell times were highly stable and conserved in human. Profiling of tRNA levels correlated with codon usage and several tRNAs were lowly aminoacylated, which was conserved in fasted mice. Finally, codons with lowly aminoacylated tRNAs and high codon usage relative to tRNA abundance exhibited long dwell times. Together, these analyses started to reveal complex dependencies between ribosome dwell times, tRNA loading, and codon usage in mammals.

## Introduction

Translation dynamically controls gene expression in processes such as development, the cell cycle, circadian rhythms, and response to stress [1]. At least three steps underlie protein translation: translation initiation, often thought to be rate limiting, elongation, and termination [2]. Recently, however, elongation has emerged as an important layer to fine-tune gene expression (reviewed in [3]). Indeed, variations in elongation rates may influence gene expression [4, 5, 6, 7, 8] and recent studies showed that alteration of ribosome elongation rates in cancer cells influences their proliferation and invasion capabilities [9, 10, 11].

While the links between translation elongation and gene expression are increasingly studied, the determinants of ribosome elongation rates are poorly understood, notably in higher eukaryotes. In unicellular organisms, ribosome elongation rates are well explained by tRNA copies and expression [12]. This is also reflected evolutionarily, since highly expressed genes are enriched for fast codons with high concentrations of tRNAs [13]. Pioneering work in *E.coli* showed that ribosomes elongation rates are different for the codons GAA and GAG [14], decoded by the same tRNA. This raises the possibility that elongation rate is not only determined by the concentration of tRNAs, but that codon-anticodon interactions as well as codon context may play important roles.

More recently, the development of ribosome profiling (RP) shed new light on the regulation of translation [15], including recently in human tissues [16]. Notably, the possibility to capture the positions of translating ribosomes on mRNAs [17] fostered the development of quantitative models providing genome-wide insights on key features regulating ribosome elongation rate [18, 19, 20, 21]. For instance, the properties of amino acids [22], (aminoacyl-) tRNA availability [23, 24, 25], tRNA modifications [26, 27, 28], secondary structures of mRNAs [29, 30, 31], folding of the nascent chain [32], pairs of codons [33, 34], and sterical interactions with the ribosome exit tunnel [35], were shown to influence the local density of ribosomes on transcripts. While RP studies have brought new knowledge on translation elongation, these were performed mostly in unicellular organisms and have led to divergent results as highlighted in several meta-analyses [19, 36]. One reason is that ribosome footprints are sensitive to biases from differences in protocols [37, 38, 39, 40, 41], library preparations [21], and data analysis pipelines [42]. Consequently, the reported correlations between elongation rates, tRNA abundances, and codon usage frequency and bias [43], show inconsistencies. In addition, while codon usage can be precisely estimated, it remains difficult to measure tRNA concentrations. Indeed, tRNAs exhibit a high degree of modifications and complex secondary structures, which alter cDNA synthesis and biases quantification by high-throughput sequencing [44]. Thus, improved methods have been proposed to quantify tRNAs [45, 46, 47, 9], as well as tRNA aminoacylation levels [48].

Here, to better establish the determinants of ribosome elongation rate in higher eukaryotes, we combined modeling of ribosome profiling data, codon usage analysis, and (aminoacyl-) tRNA profiling in mouse liver. In particular, we built a genome-wide statistical model that allowed us to identify key features predicting ribosome densities along transcripts, notably the contributions to the elongation rates of single codons, as well as pairs of codons within and near the ribosome E, P and A sites. In yeast, our analysis was able to identify novel features of previously reported pairs of adjacent codons that slow down ribosome elongation. In mouse liver, we found a wider dynamic range of codon- and amino acid-specific ribosome dwell times (DTs, defined as the inverse of the elongation rates, Methods). In addition to single-codon contributions to DTs, we identified codon pairs contributing synergistically to the DTs, and notably involving codons on the ribosome P and A sites. In mouse liver, the identified contributions of single-codon and codon-pair DTs were remarkably stable along the feeding/fasting cycle, and even under conditions of prolonged fasting. Moreover, meta-analysis in mammals revealed conserved DTs between mouse tissues and human, which were however significantly distinct from those in yeast. Finally, we extended a recent tRNA profiling method [9] to quantify (aminoacyl-) tRNA levels in liver of *ad libitum* (AL) fed and fasted mice (FA). tRNA levels correlated with codon usage and several tRNAs were lowly aminoacylated, which was conserved in fasted mice. These data, together with codon usage properties, allowed us to explain some codon-specificity in the estimated ribosome DTs.

## Results

### Modeling codon-specific DTs including single- and codon-pair contributions

Ribosome profiling read counts along transcripts typically show large variations with high and low densities of ribosomes. To estimate ribosome DTs in higher eukaryotes from those data, we explicitly fit RP read counts on a genome-wide scale, extending previous models [18, 20]. We are interested in steady-state translation, and additionally assumed no ribosome drop-off and low density of ribosomes per transcript (no traffic jams), such that the probability of finding a ribosome at a specific position on an mRNA is proportional to a position-independent gene translation flux times a position-dependent ribosome DT. To investigate how the codon context determines the DTs, we assumed that DTs depend on the codons translated in the E, P, and A sites, as well as surrounding sequences (Fig.S1A). In particular, we modeled DTs additively in log-space using single-codon contributions as well as synergistic (non-additive) contributions from pairs of codons in the E and P (noted E:P), P and A (P:A), or E and A (E:A) sites (Fig.S1A-B). The additive log-scale model for the DTs is equivalent to modeling the elongation rates at any given position with an Arrhenius rate equation, in which the activation energy has both single-codon contributions, as well as contributions from pairs of codons. Note that an inherent property of such approaches is that the DTs cannot be estimated in absolute units, but only relative to each other within each site (shown in log_2_ in the figure, Methods). Solving this model with appropriate noise distributions for RP count data can be conveniently implemented as a generalized linear model (GLM), which models the expected read counts as gene specific fluxes (gene covariates) multiplied by ribosome DTs (codon covariates) (Methods). The GLM uses the 61 sense codon alphabet, and considered positions around the ribosome spanning 40 codons around the E site (Fig.S1A-B). The same model is also applied on RNA-seq experiments (when available) to normalize the fluxes per mRNA and attenuate possible technical biases affecting the ribosome DTs. While previous models can predict ribosome footprint densities [19, 36, 30, 20, 21], our approach stands out as it allows, in addition, to determine globally translation elongation parameters (single-codon and codon-pair DTs, and gene specific fluxes) directly from the raw reads counts at every position on the coding sequences (CDSs), using negative binomial noise model (Fig.S1C-E) (Methods).

### In yeast ribosome DTs anti-correlate with codon usage and show codon-pair interactions

To validate our model, we analyzed two published ribosome profiling datasets in *Saccharomyces cerevisiae* [25, 49], one under normal (WT) [49] conditions and one treated with 3-amino-1,2,4-triazol (3-AT), which inhibits the histidine (His) biosynthesis pathway [25] thereby reducing loaded His-tRNAs. Both datasets used cycloheximide (CHX) only in the lysis buffer. Our model reproduced raw RP read counts along the transcripts with similar accuracy as previous methods [19, 36, 30, 20, 21] in both WT and 3-AT conditions (Fig. S2A-B).

Interestingly, the estimated relative dwell times (DTs) contributions for most codons in WT exhibited a two-fold range at the three E, P, and A sites (Fig. 1A-B, left), with some marked slower outliers. For instance, one codon (CCG) for proline (Pro) was markedly slow in the A and P sites, as expected due to its inefficient peptide bond formation (Discussion). Note that although the other three Pro codons are not as extreme, they are still slower than the average codon. Then, arginine (Arg) showed long DTs in all three sites. In fact, some Arg codons also showed slightly longer DTs in the upstream sequence, highlighting possible interactions of this positively charged amino acid with the ribosome exit tunnel (Fig. S2C). On the other hand, codons for isoleucine (Ile), leucine (Leu), and valine (Val) were the fastest in the A site (Fig. 1A-B, left).

**Figure 1:**
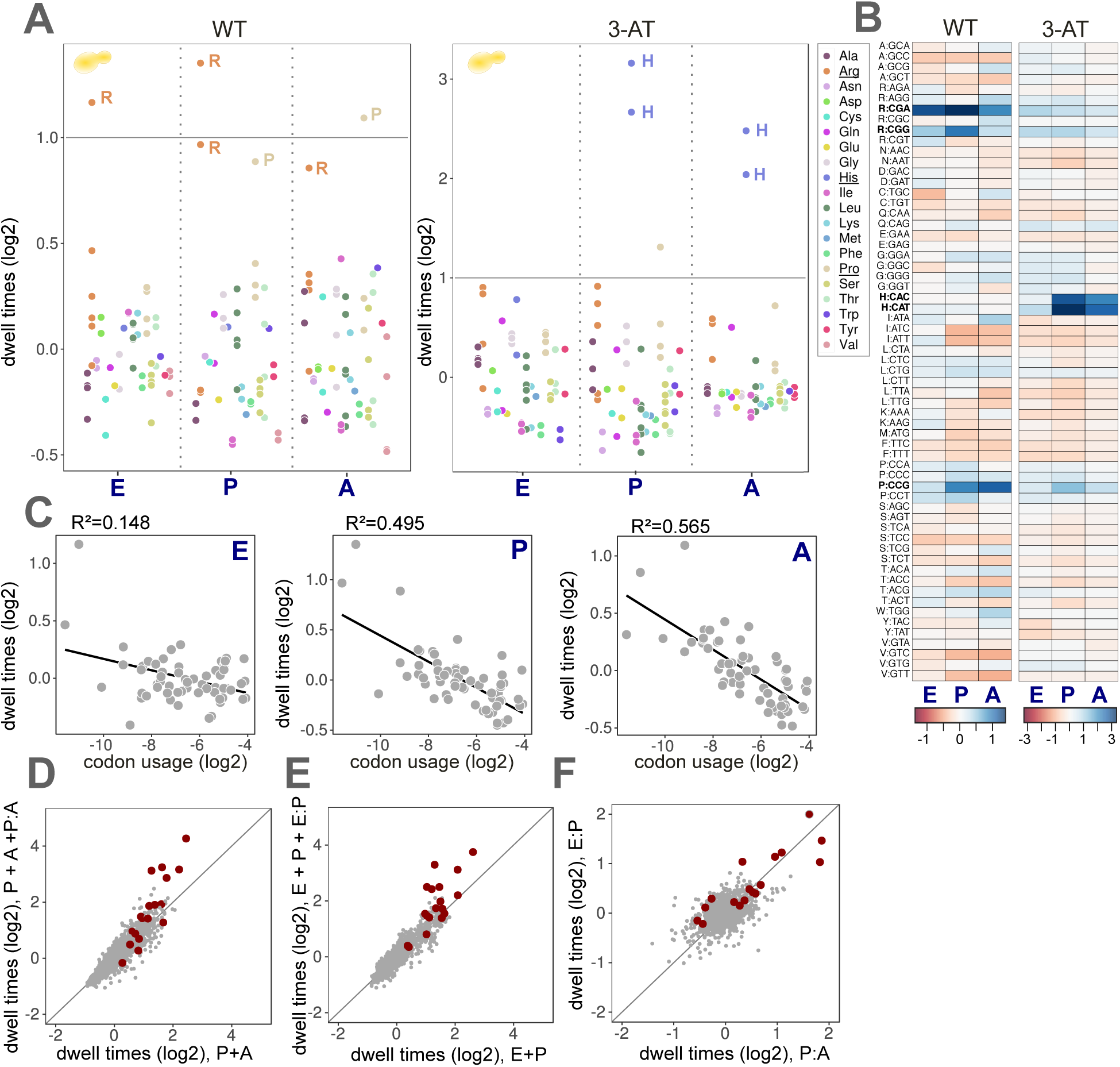
In yeast, ribosome DT anticorrelate with codon usage and show codon-pair interactions. Panels for the single-codon DTs are retrieved from the fit with the P:A interaction. A. DTs (log2, mean centered per site) for the 61 sense codons in the two conditions (WT/left, 3-AT/right) for the E, P, and A sites. Codons are colored according to amino acids. DTs with *p >*= 0.05 are not shown. Relatively fast and slow interactions are shown respectively in dark red and dark blue. B. Heatmap representation of panel (A). Here, DTs with *p >*= 0.05 are set to zero. C. Codon usage weighted by mRNA translation levels correlates with the codon DTs for the E, P, and A sites. Black line: linear fit. D. DTs (log2) for codon-pairs. Total codon-pair DTs (P+A+P:A, *i.e.* including the interactions P:A) versus the contributions from the single-codons (P+A). Red: Pairs described as inhibitory in [34]. E. DTs (log2) for codon pairs. Total codon-pair DTs (E+P+E:P, *i.e.* including the interactions E:P) versus the contributions from the single-codons (E+P). Red: Pairs described as inhibitory in [34]. F. P:A (log2) versus E:P (log2) interactions for all codon-pairs.

As control, we found that the shortage of His in the 3-AT condition resulted in lengthened DTs in the P and A sites for both His codons (CAC and CAT) (Fig. 1A-B, right). Interestingly, outside the E, P, and A sites, DTs showed a dependency on His codons at around 30 nucleotides (positions 11 and 12) downstream of the P-site (Fig. S2C), reflecting queued ribosomes (disomes) behind His codons[25]. Moreover, the DTs also displayed signatures of technical biases. Notably, the high variation in DTs at position -4, coinciding with the most 5’ nucleotide of the insert, was previously shown to reflect biases in library preparation (Fig. S2C) (reviewed in [50]). To further validate the biological relevance of our DTs, we compared ribosome DTs in WT condition with codon usage weighted by mRNA translation levels, to take into account condition-specific demands in codons. Interestingly, we found high negative correlations (*R*^2^ = 0.565 and *R*^2^ = 0.495) between the codon usage and the DTs at the P and A sites (Fig. 1C). This observation suggests an evolutionary pressure to enrich for fast codons in highly expressed genes, and conversely.

In addition to the single-codon DTs, we probed whether pairs of codons in the ribosome sites synergize by analyzing the interaction terms (E:P, P:A, and E:A) (Fig. S2D). We compared these predicted DTs with a GFP-reporter experiment in yeast probing for pairs of codons that inhibit translation [34]. Indeed, the experimentally determined inhibitory pairs exhibited long predicted DTs at the P and A, or E and P ribosome sites (Fig. 1D-E). While for these pairs the single-codon DTs were already long, the synergistic interaction terms clearly prolonged the DTs of the experimental inhibitory pairs (Fig. 1D-E). Interestingly, though the E:P and P:A interactions were not correlated overall, the inhibitory pairs stood out as showing large DT contributions in both the E:P and P:A (Fig. 1F). Globally the P:A interaction matrix was sparse and not highly structured, but revealed large values and spread for the pairs involving codons for Arg or Pro (Fig S2D). Moreover, the net DT contributions for the 61^2^ codon pairs, computed by summing the effects (in logs) of single-codon and codon pair interactions, revealed Arg in most of the top 50 slowest pairs, making this amino acid potent at decreasing translation elongation rate. Conversely, Val was contained in most of the fastest pairs. Thus, modeling RP data can identify subtle properties of ribosome DTs, such as codon-specific and codon-pairs contributions, signatures of sequences outside of the E, P, and A sites, and library biases.

### Single-codon and codon-pair DTs in mouse liver cluster by amino acids

Determinants of translation elongation are less studied in mammals. We and others have previously shown that feeding/fasting cycles can regulate translation initiation in mouse liver [51], via well described mechanisms, notably through mTOR and GCN2 related nutrient sensor pathways (reviewed in [52]). Here, we aimed to extend this analysis to the level of DTs, in particular to asses whether perturbed amino acid pools during low nutrient availability can alter DTs. Therefore, we applied the above model to our previously 84 ribosome profiling samples harvested in WT and circadian clock deficient mice (*Bmal1* KO) every 2-4 h around the 24 h day, including four biological replicates. As for yeast, our model faithfully captured raw RP read counts along transcripts, as reflected by a previously used correlation metric (Fig. S3A-B). Remarkably, DTs were very stable across all samples, showing high biological reproducibility and no time or circadian clock-dependent changes (Fig. S3C-E). Therefore, for the following analyses, we averaged the DTs over all the 84 samples.

Patterns of DTs for the E, P, and A sites showed biologically significant codon and amino acid-specificity (Fig.S3C). DTs were strikingly different than in yeast and exhibited a larger dynamic range (Fig. 2A-B). In particular, the P and A sites revealed nearly 10-fold range in DTs between the fastest and slowest codons, while DTs in the E site were more tightly distributed (Fig. 2A-B), presumably reflecting that the DTs are primarily sensitive to aminoacyl-tRNA availability (A site) and peptide bond formation (P-A sites). While DTs in the P and A site were overall more strongly correlated to each other than with the E site, DTs also showed clear site-specificity (Fig. 2A-B). For instance, the four codons for Gly had long DTs in the P site, however, the Gly GGT codon was among the fastest in the E and A sites, while the GGA codon was markedly slow in the A site. Strikingly, all three Ile codons had long DTs in the A site but very short DTs in the P site (Fig. 2A-B). For the negatively charged glutamate (Glu) and aspartate (Asp), all their codons showed long DTs in the P and A sites (Fig. 2A-B). Considering a larger window around the ribosome revealed that P and A sites, followed by the E site, showed the largest contributions to DTs (Fig. S4A). Upstream and downstream sequences outside the (−4,+6) interval did not contribute (Fig. S4A), while codons in the vicinity of the ribosome (−3, −2, −1 and +3, +4, +5) exhibited significant variations in DTs, and were correlated between both sides. The detected signals at the −4 and +6 positions reflect known ligation biases during the library preparation [50].

**Figure 2:**
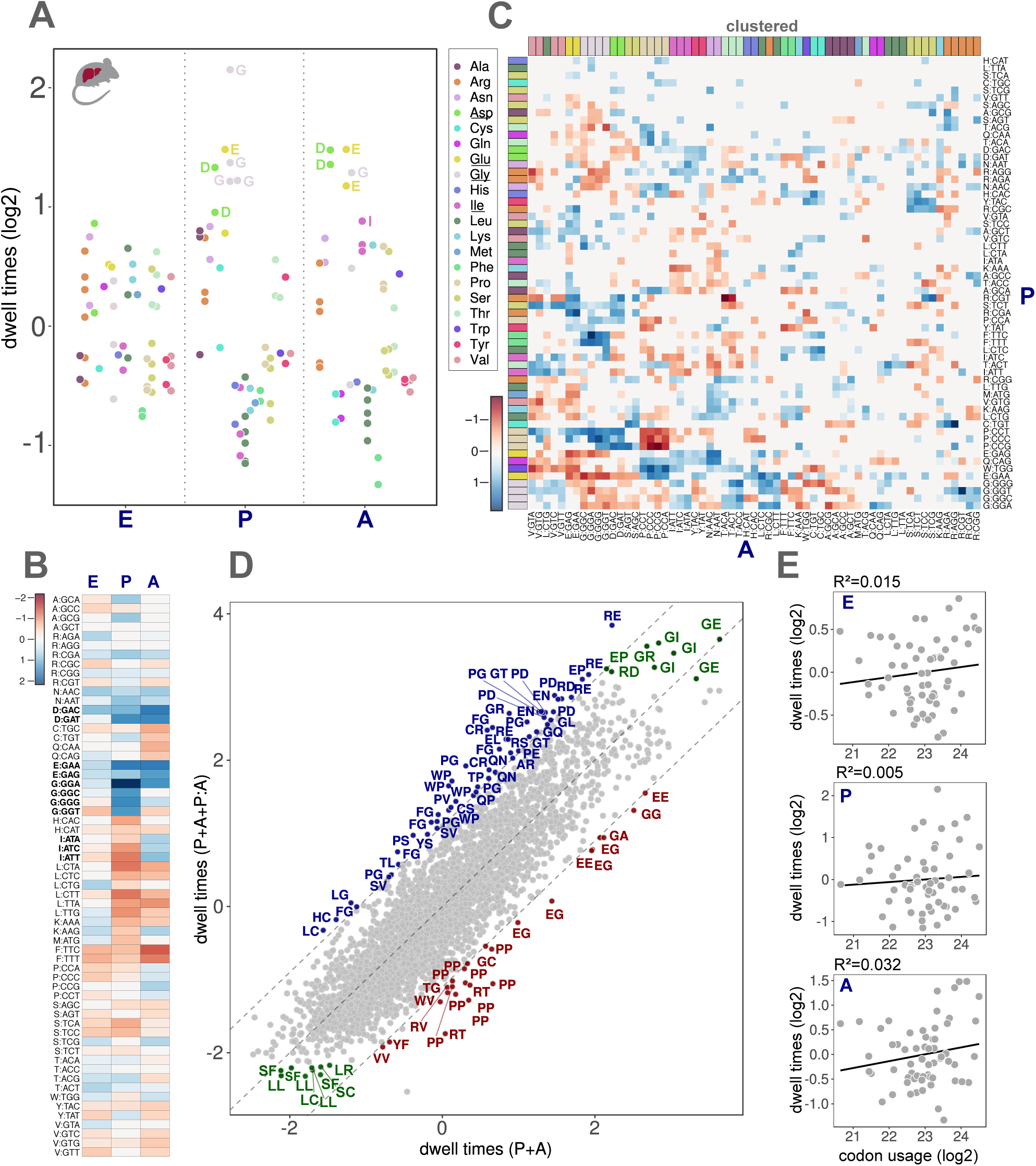
Single-codon and codon-pair DTs in mouse liver cluster by amino acid. Single-codon DTs are retrieved from the fit with the P:A interaction. A. DTs (log2, mean centered per site) for the E, P, and A sites averaged over the 84 samples in mouse liver. Codons are colored according to amino acids. DTs with *p >*= 0.05 are set to zero. Fast and slow DTs (relative to the mean) are shown in dark red and dark blue, respectively. B. Heatmap representation of panel A). DTs with *p >*= 0.05 are set to zero. C. Interaction matrix for the pairs P:A (log2). Codons are colored according to amino acids. Codons in both sites are hierarchically clustered (euclidean distance matrix, complete linkage algorithm). Fast and slow interactions are shown in dark red and dark blue, respectively (colorbar). D. DTs (log2) for codon pairs. Total codon-pair DTs (P+A+P:A, *i.e.* including the interactions P:A) vs. the contributions from the single-codons (P+A). Pairs with interactions *>* 1.1 or *<* − 1.1 are annotated and colored, respectively, in blue and red. Top 10 slowest and fastest pairs are colored in green if not already depicted. E. Codon usage weighted by mRNA translation levels does not correlate with codon DTs for the E, P, and A sites. Black line: linear fit.

Codon-pair DTs revealed a significant influence on translation elongation in mouse liver, with P:A interactions showing the widest dynamic range, followed by E:P and E:A (Fig. S4C). Note that the interaction matrices are not symmetric, showing codons or amino acid specificity at the respective ribosome sites (Fig. 2C and S4B). Intriguingly, the P:A matrix highlighted a striking clustering by amino acid for the A site (Fig. 2C), while E:P interactions clustered by amino acid in the P site (Fig. S4B). This suggests that P:A codon-pair DTs are determined by amino acids through their influence on the peptide bond formation. The clustering by amino acid was corroborated by a model selection analysis on the 84 samples, where the alphabet for the DT regression coefficients was taken as either the 20 natural amino acids, or the 61 sense codons (Fig. S4D). While the preferred alphabet was overall that of the codons, the model with amino acid coefficients at the A site for the P:A interaction was preferred to all the other models (Fig. S4D). In the case of the E:P interaction, the amino acid alphabet in the P site was considered as the best model. Overall, models including the site interactions were preferred to the reduced models, emphasizing the importance of codon-pair interactions in determining ribosome DTs in mouse liver. The P:A matrix revealed strong positive interactions (ones lengthening the DTs) for pairs of bulky (Pro, tryptophan (Trp) and phenylalanine (Phe)) or achiral (Gly) (Fig. 2C and D) amino acids. As in yeast, Arg codons were frequent among large P:A contributions (Fig. 2C and D). Surprisingly, the known stalling pair Pro-Pro showed the largest negative interaction (Fig. 2C and D), possibly related to eIF5A activity which is known to facilitate otherwise slow peptide bond formation for such pairs [53]. Overall, these interactions contributed to the total codon-pair DTs for the P and A site by a factor larger than 2 for about one hundred pairs (Fig. 2D). Summing both single-codon and codon-pair contributions showed that the amino acid pair isoleucine-glycine was represented by multiple combinations of codons in the top ten overall slowest pairs (Fig. 2D). Similarly the leucine-leucine pair was frequent among the fastest codon pairs (Fig. 2D). The E:P matrix was more complex: pairs involving the amino acids Gly, Asp, Asparagine (Asn), and Pro in the P site lengthened the total codon-pair DTs (Fig. S4B). Unlike in yeast (Fig. 1C), ribosome DTs did not correlate with codon usage in mouse liver (Fig. 2E), arguing for different evolutionary pressure on translation efficiency.

### Ribosome DT patterns in liver are stable under prolonged fasting

The above analysis showed highly robust DTs between liver samples collected during the normal feeding (night) / fasting (day) cycle (Figure S3C-E). To probe whether ribosome DTs are sensitive to longer periods of fasting, we performed new ribosome profiling experiments in mice fed either ad libitum (AL) or fasted (FA) for up to 30 hours (Fig. 3A). Since the enrichment in ribosome footprints can be sensitive to RP protocols [54], we here used a small RNA-Seq protocol with random adapters (UMI) to reduce possible ligation biases and PCR duplicates. Moreover, as ribosome dynamics and DTs are affected in ribosome profiling experiments with cycloheximide (CHX) in yeast [39, 55], we tested conditions without CHX in the lysis buffer. First, we validated the effect of prolonged fasting by analyzing differential ribosome profiling signals between AL and FA. Genes related to the Peroxisome Proliferator-Activated Receptor *α* (PPAR*α*) pathway and to fatty acids oxidation were upregulated in FA, presumably to provide the energy needs (Fig. 3D). On the contrary, genes related to lipid biosynthesis were downregulated in FA (Fig. 3D), suggesting that animals switched from glucose to fatty acid metabolism in FA, as already described [56]. Moreover, *Mat1a*, *Asl* and *Got1* related to amino acid biosynthesis were upregulated in FA (Fig. 3D), presumably in response to perturbed amino acid homeostasis. The ribosome profiling data in the new conditions showed a typical tri-repeat nucleotide pattern (Fig. 3B), confirming the presence of *bona fide* translating ribosomes in the FA samples, as well as in samples without CHX (NOCHX). In addition, the modified library preparation improved the goodness of the fits. Indeed, correlation coefficients introduced above (Fig. S3A-B) were now improved (Fig. S5A-B).

**Figure 3:**
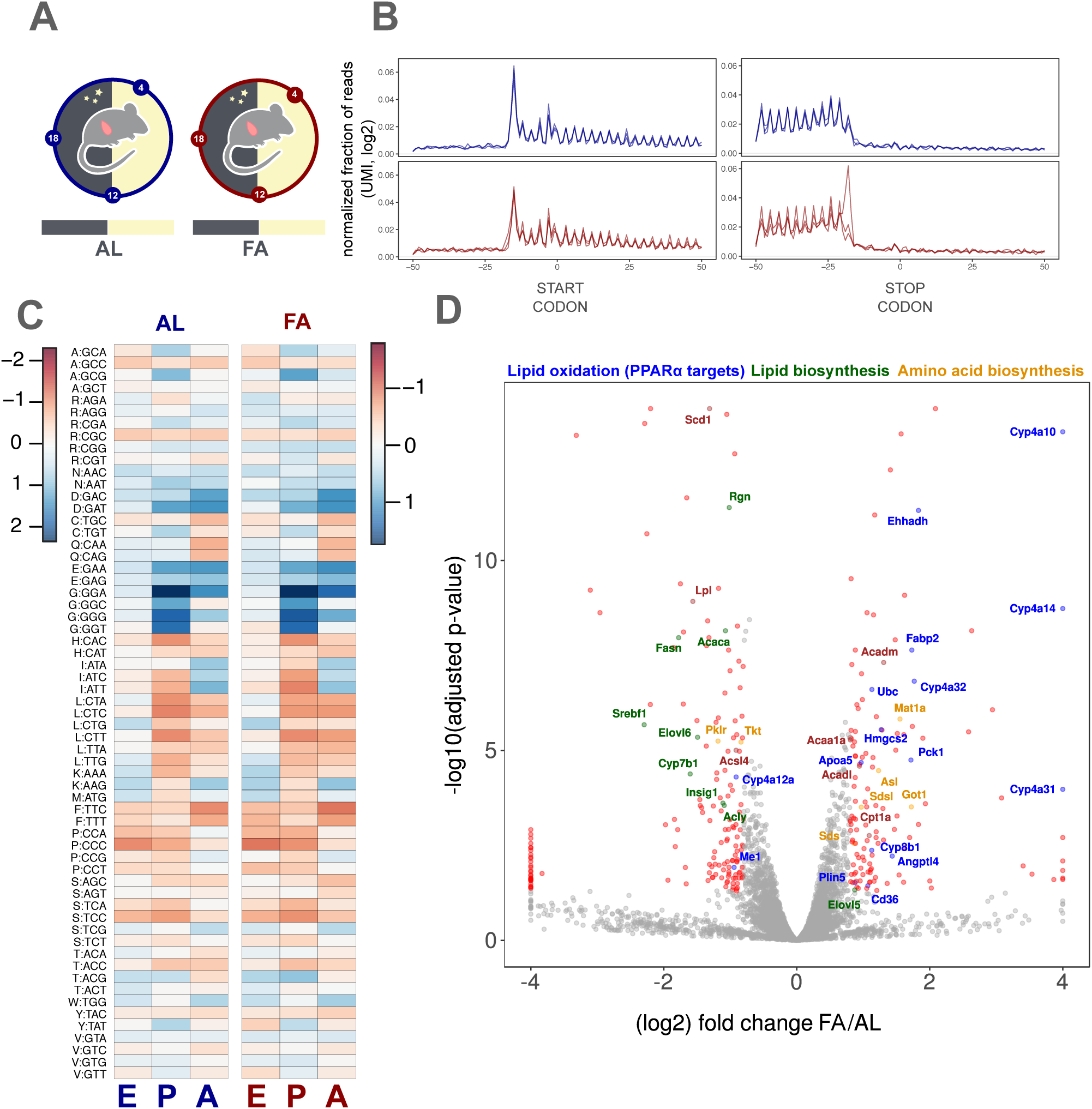
Ribosome DTs are not affected in fasted mice and without cycloheximide. Single-codon DTs are retrieved from the fit with the P:A interaction. A. Livers from mice fed *AL* or FA were harvested at ZT4, ZT12, and ZT18. Ribosome profiling was performed without cycloheximide (CHX) in the lysis buffer. B. Normalized fractions of reads of length 32 around the start and stop codons in a window of 100 nucleotides genome-wide. Dark blue: AL/NOCHX; dark red: FA/NOCHX. C. DTs (log2, mean centered per site, heatmap) for the E, P, and A sites in AL/NOCHX and FA/NOCHX. Codons are ordered by amino acid. Side bars: log2 color scale. D. Differential expression of ribosome profiling signals between AL and FA (Methods). Benjamini-Hochberg adjusted p-values (-log10) plotted against averaged log2 fold change between FA and AL. Genes with false discovery rate (FDR) *<* 1% and absolute log2 fold change *>* 1 are annotated and colored. Blue: genes in the KEGG “PPAR*α* signal pathway”; Green: KEGG and GO term “lipid biosynthesis”; Orange: KEGG “amino acid biosynthesis”.

Surprisingly, single-codon DTs in the AL and FA conditions showed no difference in both the CHX or NOCHX conditions (Figs. 3C and S5C), and codon-pair DTs very also very similar (Fig. S5D). Moreover, these DTs were highly correlated with those described above for the 84 samples (Fig. S6C-D). Nevertheless, the FA samples showed a reduced dynamic range in the DTs, presumably due to variability in ribosome profiling signal quality (Fig. S5A) related to the global decrease in translation levels.

Finally, we probed whether the perturbed metabolic state in FA might lead to differential codon usage [24]. Strikingly, when considering the codon usage bias in WT and FA animals, we found that most of the codons with a G or C nucleotide at the third position (GC3) were enriched in up-regulated transcripts in FA, while codons with an A or T nucleotide were underrepresented (Fig. S5E).

### Meta-analysis reveals conserved ribosome DTs in mammals

To further compare the estimated ribosome DTs, we analyzed published ribosome profiling datasets in mouse liver (H: Howard *et al.*) [57], mouse kidney (CS: Castelo-Szekely *et al.*) [58], and in the human liver cell line Huh7 (L: Lintner *et al.*) [59]. Despite differences in library preparation protocols (Methods), single-codon DTs at the A site were highly correlated between the mammalian datasets (0.48 *< r <* 0.96), including in different tissues (kidney) and human cells (Fig. S6C, E). However, the mammalian DTs were markedly different from those in yeast [19]. Note that the relative contribution to the DTs from the E, P, and A sites vs. that from surrounding sites, in particular at positions −4 and +6, differed significantly depending on the protocols (Fig. S6A-B). Similar potential biases have been reported previously [54]. Here, we found that protocols without cDNA circularization showed highest signals in the P and A sites, presumably reflecting ribosome dynamics more faithfully (Fig. S6A-B). Moreover, this was also reflected in the codon-pair DTs (P:A), which were more consistent across experiments with that protocol (Fig. S6D).

Together, this meta-analysis highlighted how different library preparations lead to damped RP signals in the A and P sites for some protocols, and showed that the codon DT patterns are conserved between mouse tissues and mammalian species.

### (Aminoacyl-) tRNA profiles are conserved in fed and fasted mice

We next asked whether the estimated DTs can be linked with tRNA abundances or loading levels, which is poorly studied in higher eukaryotes [43]. The chemical modifications and secondary structure of tRNAs render them difficult to quantify. A recent hybridization method combined with sequencing, which controls specificity using left/right probes and a stringent ligation step, allows to bypass the cDNA synthesis to quantify tRNA levels [9]. To measure tRNA abundances and assess possible links with ribosome DTs in mouse liver, we adapted and optimized this method to target all annotated mouse tRNAs (Fig. S7A). Moreover, we quantified the fraction of (aminoacyl-) tRNAs using sodium periodate [60], which depletes unloaded tRNAs by selective biotinylation of 3’-ends (Fig. S7A). This way, we aimed to quantify the tRNA pools available for elongation in the ribosome A site.

tRNA molecules are encoded by a large number of genes. Therefore, we designed 303 DNA probe pairs (left and right) to target all mouse tRNA sequences from the *GtRNAdb* database [61] (Fig. S7A). Our modified protocol yielded a high proportion of specific ligations between left and right probes, showing target specificity for tRNAs (Fig. S7B). Indeed, mapping of the sequencing reads to all possible combinations (303^2^) of left and right probes showed that more than 75% of ligated products belonged to tRNA genes of the same codon (Fig. S7B), and even 95% were from probe pairs that could be assigned to specific codons with high confidence (Fig. S7B) (Methods). We performed further experiments to validate the specificity and evaluated the efficiency of DNA ligases (Fig. S7C-D).

We measured tRNA abundance on mouse livers from the same samples as those used for the ribosome profiling. Specifically, we quantified the total tRNA (control, NaCl) and the (aminoacyl-) tRNA (sodium periodate, NaIO_4_) abundances from the same pieces of liver in two replicates in the AL and FA conditions, at three different times during the day (ZT04, ZT12, ZT18) (Fig. 4A, Methods). tRNA abundances were highly reproducible (Fig. S7E), exhibited a large dynamic range (Fig. 4B), and were significantly correlated with PolIII ChIP-Seq data in mouse liver [62] (Fig. S7F). tRNA levels for amino acids encoded by four synonymous codons (“4-codon” box) were represented by one dominant highly expressed isoacceptor with a T at the wobble position 34 (*e.g.* TGC/Ala, TGG/Pro, TCC/Gly, TAC/Val, TGT/Thr) (Fig. S8A). The tRNA levels showed only small variations over the biological conditions, except for mitochondrial tRNAs (Fig. S8B). Strikingly though, principal component analysis (PCA) on the four conditions (*i.e.* NaCl/AL, NaCl/FA, NaIO_4_/AL and NaIO_4_/FA) showed a clear separation between the control and periodate conditions (Fig. 4C), indicating differential loading of the tRNAs. Surprisingly, the AL and FA samples were indistinguishable in the total tRNA and (aminoacyl-) tRNA conditions (Fig. 4C-D), indicating no imbalance of tRNA charging in prolonged fasting and coinciding with the stable DTs.

**Figure 4:**
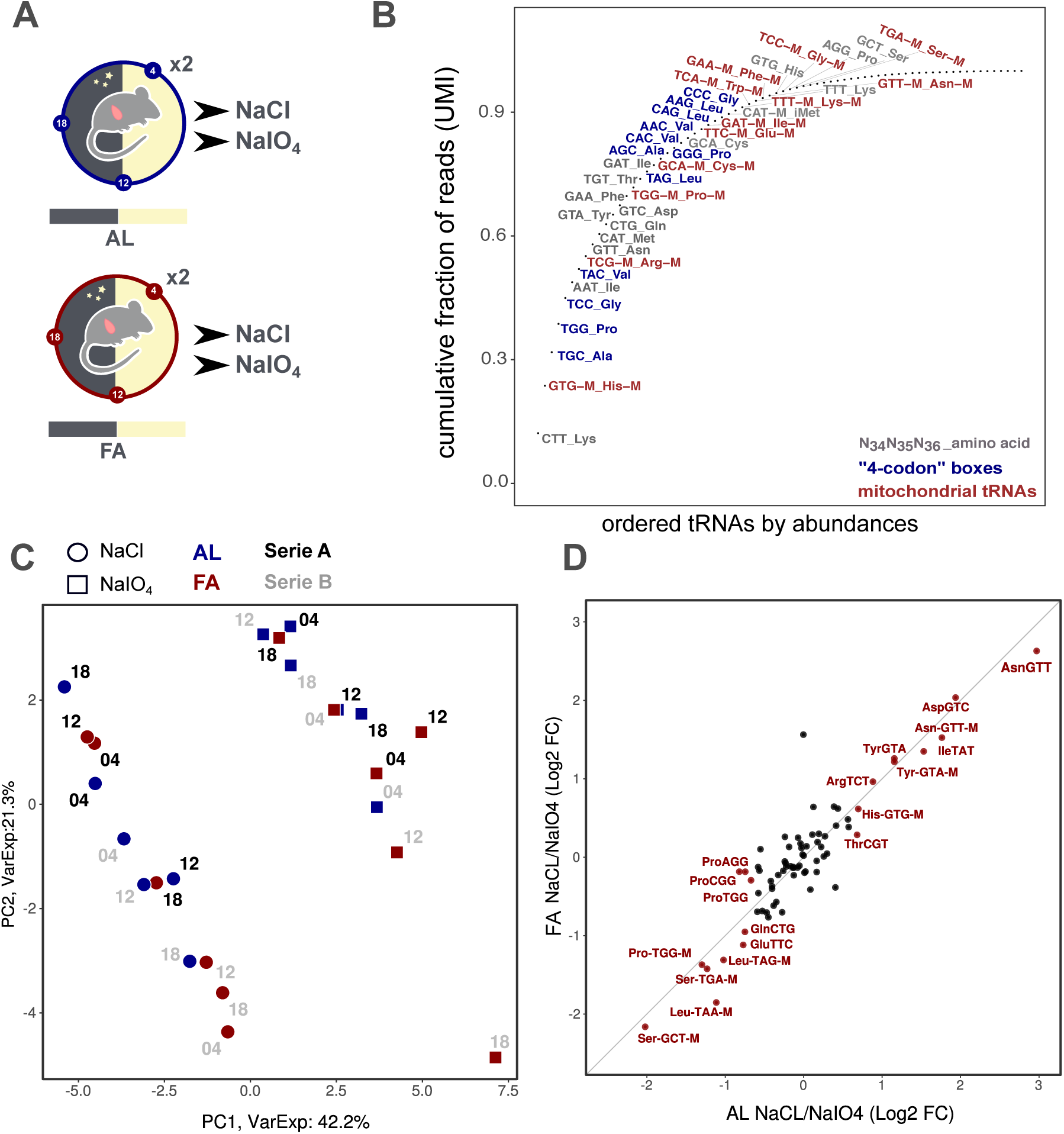
(Aminoacyl-) tRNA profiling in AL fed and FA mice. A. Mice fed AL or FA were sacrificed at ZT4 (fasting duration: 16 h), ZT12 (24 h) and ZT18 (30 h) and livers were harvested. Each sample was treated with NaCl and sodium periodate (NaIO_4_). B. Cumulative fraction of reads for each tRNA, ordered by abundances. Anticodons and amino acids are indicated for the 50 first codons. Blue: four-codon box amino acid; red: mitochondrial tRNAs. C. PCA of the tRNA abundances (log2 UMI). PC1 and PC2 explain 42.2% and 21.3% of the variance, respectively, and separate NaCl from NaIO_4_ treatment. NaCl (circle), NaIO_4_ (square), AL (blue), FA (red), replicate 1 (black), replicate 2 (grey). ZT is shown beside the points. D. Ratio of tRNA abundances (log2 fold change, averaged over the time points) between the NaCl and NaIO_4_ for AL fed vs. FA mice (significant changes, *p <* 0.05 in red). No tRNA showed a significant difference between AL and FA (*i.e.* fell out of the diagonal).

However, some codons for Asn, Asp, and Ile were lowly aminoacylated, independently of the feeding regime (Figs. 4D, S8C). Strikingly, these same codons were found to exhibit the slowest DTs in the A site (Fig. 2).

### Relationship between (aminoacyl-) tRNA levels, codon usage, and dwell times

To substantiate this observation, we investigated whether variations of single-codon DTs and codon usage could be explained by the available tRNA pools. Here, our data showed a significant correlation between codon usage and our measured tRNA levels in mouse liver (Fig. 5A), extending previous work using POLIII loadings on tRNA genes as proxies [63]. Our analysis also highlighted codons with high or low demand (codon usage) compared to the supply (tRNA levels), as quantified by the *codon balance* [64] (Fig. 5A).

**Figure 5:**
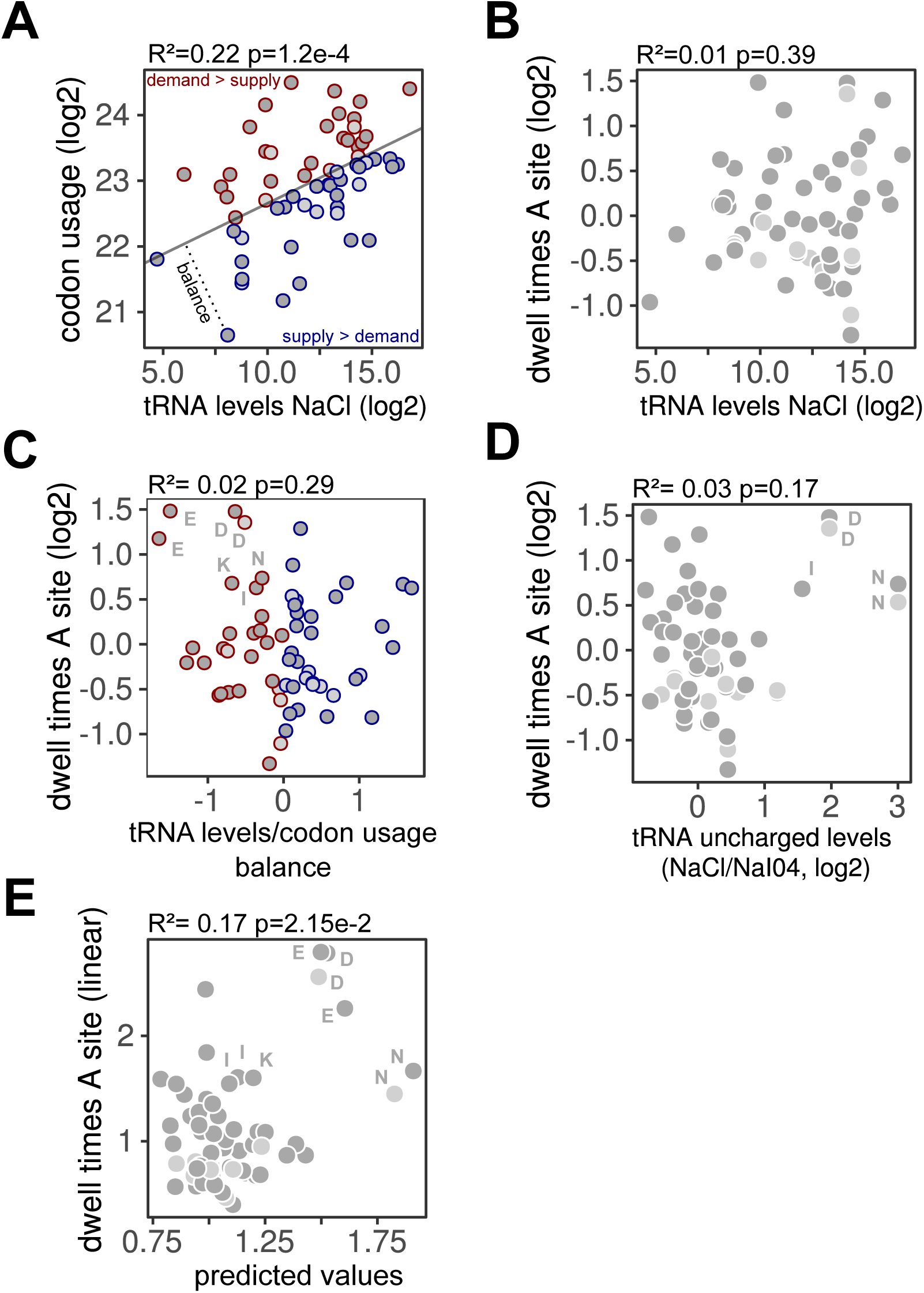
Relationship between (aminoacyl-) tRNA levels, codon usage and DT. A. Significant correlation (*R*^2^ = 0.22*, p* = 1.2*e*−4) between codon usage weighted by mRNA translation levels (wCU, log2) and normalized total tRNA (NaCl) read count for each codon (log2, UMI, averaged over the AL samples). The gray line shows the first principal component (PC). Orthogonal distance to the PC reflects the balance between tRNA supply and demand. Codons with positive (resp. negative) balance are colored blue (resp. red). Codons are assigned to their canonical tRNAs (dark-grey) or to wobble tRNAs (light-grey) where appropriate (Methods). B. DTs in the A site (log2) *vs.* tRNA levels (NaCl, log2) averaged over the AL samples (*R*^2^ = 0.01, p-value= 3.9*e*−1). C. Correlation between tRNA levels/codon usage balance and DTs in the A site (log2) averaged over the AL samples. *R*^2^ and p-value are reported for the linear regression (*R*^2^ = 0.02,p-value= 2.9*e*−1). D. Correlation between tRNA uncharged levels (NaCl/*N aI*0_4_, log2) and DTs in the A site (log2) averaged over the AL samples. *R*^2^ and p-value are reported for the linear regression (*R*^2^ = 0.03, p-value= 1.7*e*−1). Positively correlated codons are annotated by their one-letter amino acid. E. Significant correlation(*R*^2^ = 0.17, p-value=2.15*e*−02) between estimated and predicted DTs in the A site. Prediction uses a linear model with the balance and uncharged levels, in linear scale, as explanatory variables. In C-E, annotations refer to one-letter amino acid.

Some codons with the slowest DTs in the A site clearly stood out as having either low codon balance or aminoacylation levels. In fact, DTs in the A site did not exhibit a simple correlation with tRNA abundances (Fig. 5B), nor with the codon balance (Fig. 5C). However, the slow DTs for Glu codons (Fig. 5C) may well result from their low codon balance, hence limiting tRNA availability at the A site. Remarkably, codons for Asp, Asn, and Ile, which had particularly lowly charged tRNAs, coincided with some of the slowest DTs in the A site (Fig. 5D). However, DTs in the A site were overall uncorrelated with tRNA aminoacylation levels. We therefore included several effects in a linear model, which uncovered that a linear combination of tRNA aminoacylation levels and codon balance captures a significant portion of variation in the A site DTs, particularly the long DTs for Glu, Asp, Asn, and Ile codons (Fig. 5E).

## Discussion

We extensively modeled RP datasets and uncovered single-codon and codon-pair DTs determining ribosome elongation rates in mammals. These DTs were highly stable across all conditions tested. In parallel, we quantified (aminoacyl-) tRNA levels in mouse liver and identified several features regulating ribosome elongation, such as aminoacylation levels and the balance between tRNA levels and codon usage.

In yeast, our model accurately inferred codon-specific DTs and highlighted mainly Arg and Pro as slow in the A and P sites. These amino acids are known for their inefficient peptide formation and sterical interactions with the ribosome [65, 66]. A significant negative correlation was observed between DTs and codon usage, reflecting natural selection for fast codons in highly translated genes. While this relationship has been described [67, 36], the found correlation is, to our knowledge, the highest reported. Moreover, our analysis confirmed recently identified inhibitory pairs [34], and deciphered their synergistic effect in addition to the site-specific contributions. We showed that the inhibitory pairs lengthened DTs both in the E:P and P:A positions, highlighting potentially inefficient translocation of the pair due to wobble base-pairing or other mechanisms [34].

In mouse liver, DTs differed significantly from yeast, showing a larger spread and higher complexity. Remarkably, DTs were very similar between different tissues, species, and RP protocols. Moreover, the DTs were consistent with a peptide motif enriched in stalled ribosome sites in mouse embryonic stem cells (mESCs) [68].

We found that the smallest and achiral amino acid Gly exhibited very long DTs (in the A and P site) with different magnitudes between the isoacceptors and tissues (*i.e.* liver and kidney). Interestingly, in bacteria, Gly codons are slow, although this effect is still difficult to separate from Shine-Dalgarno (SD) dependent stalling [69] or protocol artifacts [70], and is therefore debated [71]. As mammals do not use a SD mechanism, our result support an alternative hypothesis, such as slow codon-anticodon pairing [72] or inefficient peptide bond formation. Pioneering work in *E. coli* suggested that Gly-tRNAs adopt a particular conformation due to the U nucleotide in position 32 and that unmodified *U*_34_ on 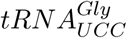 could decode the four Gly codons (a pairing known as superwobbling [73]), but with low efficiency (reviewed in [72]). While this mechanism was shown in unicellular organisms, our tRNA profiling found 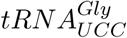 as the major Gly isoacceptors and one of the most abundant tRNAs in mouse liver. Further work may indicate whether superwobbling occurs in the liver.

The DTs for the acidic amino acids (Asp and Glu) were among the slowest. Glu showed a particularly low balance of tRNA levels/codon usage and Asp tRNA was lowly charged. This could lead to a shortage of tRNA availability and ribosome stalling [23]. As their codons share the same first two bases, competition with near-cognate tRNAs [27], or pairing inefficiency due to the wobble mechanism could also explain the long DTs. Indeed, slower elongation would allow higher precision in codon-anticodon discrimination [74].

Ile codon DTs were slow in the A site while fast in the P site. Remarkably, the isomeric Leu codons were the fastest in the P and A sites, highlighting a structure-independent mechanism. Indeed, we showed that Ile-tRNAs were lowly aminoacylated, reducing Ile availability on the A site, but other explanations are possible. For instance, since Ile is decoded by three different codons, a suitable pairing mechanism such as inosine or other U34 modifications could be used to avoid pairing of the fourth near-cognate codon (Met) and therefore increase the DTs [75].

One of our main findings concerned the contributions of codon-pair interactions towards DTs, mainly at the P and A site. At these positions, the ribosome catalyzes the peptide bond formation between (aminoacyl-) tRNA in the A site and peptide-tRNA bound to the P site. Our analysis revealed that the identity of the amino acid in the A site (acceptor), and not the codon, was the best descriptor of those codon pair interactions. Pairs including bulky amino acids or Gly in the A site were slow, highlighting their potential inefficiency in peptide bond formation. Interestingly the DT for Pro-Pro pairs, known to inefficiently form peptide bonds [76], was markedly reduced by the interaction. This observation probably shows the role of eIF5A in resolving this stalling motif. On the other hand, Gly, Asp, and Glu, which were slow in our analysis, were shown by others to require eIF5A for their efficient translation [77, 78].

Other features not included in the model, and which are independent of the codon identity, might regulate ribosome elongation. A high number of liver proteins are secreted and thereby translated by ribosomes bound to the endoplasmic reticulum via the interaction of signal recognition particles with the nascent peptide chain. These interactions are known to stall the ribosomes, however, as these appear to be codon independent, we did not detect them in our analysis [30]. In addition, chaperone proteins interacting with the nascent peptide can influence co-translational folding and subsequent ribosome density on mRNAs [79]. The model could also be further extended by including RNA secondary structure and modifications, pseudo-knots acting as ribosome roadblocks, and slippery sequences inducing frame shifting [30, 20, 80, 81]. Recent studies have described ribosome collisions and their relationships with recruitment of ribosome quality control and degradation pathways [82, 83]. While these events could happen frequently in liver, and thereby bias position-dependent estimation of DTs from standard ribosome profiling, a recent study probing the determinants of collided ribosomes in mouse liver showed similar codon dependencies between pausing sites and our DTs [84].

While we found a striking correlation between DTs and codon usage in yeast, the same did not hold in mammals. This suggests that biased codon usage in mammals reflects more complex evolutionary forces, such as mutation driven GC bias [63]. Nevertheless, the measured tRNA abundances showed signatures of adaptation, since tRNA levels correlated with the codon usage. These correlations extended previous results at the transcription level or in highly expressed genes [63, 85]. Related to this, one still open challenge is to assign tRNAs to their corresponding codons, due to the extended wobble base pairing rules related to tRNA modifications.

Surprisingly, tRNA loading was unaffected by prolonged fasting. Several studies in cell lines showed that decreasing amino acids in culture media leads to decreased (aminoacyl-) tRNA availability and therefore increases ribosome stalling [60, 23]. Moreover, others have shown that codon optimality contributes to differential mRNA translation in response to starvation [24]. While we did not observe this, probably due to the *in vivo* state, GC3 bias (*i.e.* GC bias at position N3 in codons) was significantly different between genes translated in AL and FA mice (Fig. S5E) or also between night and day conditions (not shown). Genes with high GC3 content have been shown to provide more targets for methylation than those with low GC3 and to be enriched in stress responsive genes [86]. Oxidative stress occurs during fasting/day in mouse and correlates with GC3 content. Nevertheless, the reason of the higher GC3 level in FA compared to AL still needs to be identified.

Like (aminoacyl-) tRNA levels, DTs were unchanged between AL and FA. We can hypothesize that after more than 30 hours of starvation, mice compensate the lack of amino acids by a large global decrease of translation initiation through mTORC1/GCN2 [87], making tRNA availability and translation elongation non limiting. Moreover, since ribosome profiling signals, DTs, and tRNAs were measured in relative and not absolute amounts, we cannot exclude a total decrease of translation elongation rate, aminoacylation, or tRNA levels.

In conclusion, ribosome DTs, codon usage, tRNA levels, and translation elongation in mammals do not obey simple relationships. Nevertheless, although a global understanding is still missing, we were able to link both tRNA/codon usage balance and aminoacylation levels with anomalously slow DTs in the P and A site of the ribosome. Probing different ribosome states (*e.g.* free A site) using RP combined with different drugs [70] or improving the quantification of (aminoacyl-) tRNA through nucleotide modification removal [45] will lead to better understanding of the determinants of translation elongation. Finally, more work is needed to understand the consequences of changes in ribosome elongation rates for mRNA stability and nascent protein folding.

## Methods

### Inference of dwell times and translation fluxes

#### Preprocessing of ribosome profiling data

RP datasets from yeast, mouse, and human were respectively mapped on the sacCer3, mm10 and Hg38 genomes using STAR [88] with parameters –`seedSearchStartLmax 15`. Genomes indexes were built using Ensembl transcripts annotations. Adapters were retrieved for the different datasets and input as parameter for STAR. In the case of NEXTFlex library, fastqs files were parsed and duplicated sequences (UMI and insert) are removed. Sequences were trimmed for adapters using fastx_clipper with parameters `-Q33 -a TGGAATTCTCGGGTGCCAAGG -l 11` and UMIs are removed (4 nucleotides on both sides). Then, the fastq files are mapped using STAR with options –`seedSearchStartLmax 15`. The subsequence BAM files were sorted and indexed.

#### Read counting on the coding sequences

For each protein coding transcript with a CDS larger than 120 nucleotides, reads with zero mismatches, unique mapping (nM:i:0 & NH:i:1) and a length between 25 and 40 nucleotides were retrieved using `samtools view` in the respective region. E site position was defined, for each read size, in function of the frame on the CDS and pileup plots at the start codon. From the E site position, the sequence in the window [-60,+60] nts was reported and incremented by one at each new observation. Sequences with a window spanning the start or stop codon were removed.

#### Data filtering

A sliding window of 120 nucleotides moving 3 by 3 on the CDS of protein coding genes were computed and the respective sequences were reported (Figure S1D). This set of sequences is used as a reference and their respective number of counts is set to zero. Every time a read occurs at one of these sequences, we incremented the count by one (Fig.S1D). Genes with less than 5% of positions covered or less than 5 positions observed were discarded. Genes with less than 100 counts were removed. Sequences containing a stop codon (TAG, TGA, or TAA) or non-unique in the coding genome were discarded. Depending on the sample coverage, we monitored about 5000 genes in mammals.

#### Generalized linear model for ribosome profiling count data

We used a generalized linear model for the observed RP read counts at the different positions on the gene CDS. Here, the read counts *Y_igs_* at a specific codon position *i* corresponding to the ribosome E site on the CDS of a gene *g* in sample *s* were modeled as a negative binomial (NB) (*i.e.* commonly used for overdispersed count data [89]) with mean *µ_igs_* and dispersion parameter *θ_s_*.

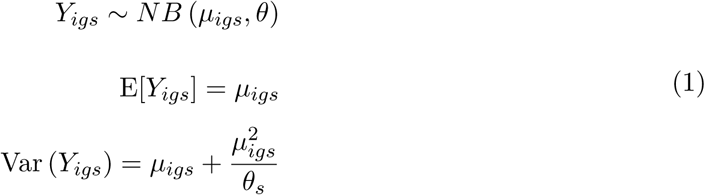

The mean is further modeled as follows (omitting the sample index) :

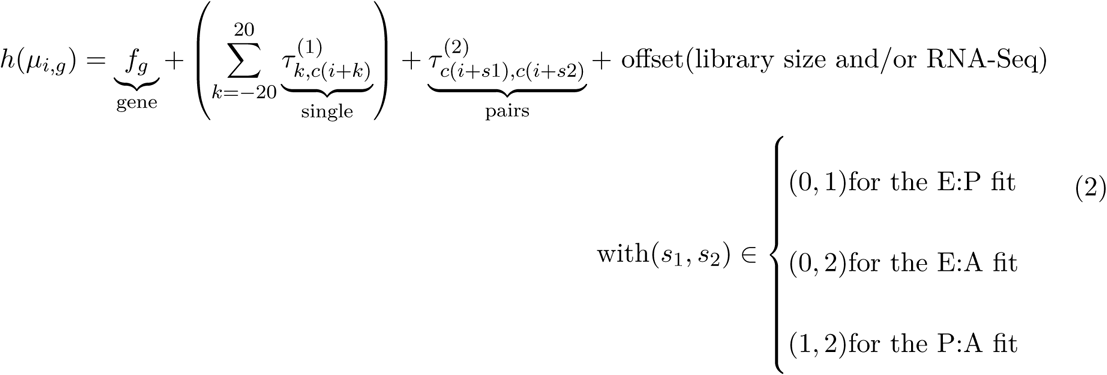

Here, *c*(*i*) *∈ {AAA, AAC, AAG, …, T T T }*. *h*(*x*) = log(*x*) the so-called link function that allows to express the product of gene flux and DTs (equal to the expected RP density) as a sum in log-space. 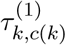 denote the contributions of single-codon DTs (in log scale) for the 61 sense codons at position *k*, with *k* the position relative to the ribosome E site. 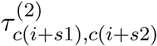 denote the contributions of codon pair DTs (again in log scale) for the 61^2^ pairs of sense codons at positions (*s*_1_*, s*_2_) relative to the ribosome E site. These codon pair matrices are modeled for the sites E:A, E:P, and P:A. *f_g_* is the gene-specific translation flux (in log scale). Note that since this problem does not have full rank, we must choose some constraints, which means that the fit DTs are *relative* contributions. Here we chose to express these contributions relative to the mean for each site. Specifically, we apply the convention: 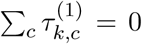 for all *k*, 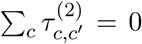 for all *c*′, and 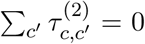 for all *c* and shifted the gene fluxes accordingly. The offset term(s) make the gene fluxes normalized by library size. In addition, when specified in the text, we normalized gene translation fluxes by mRNA abundance by fitting the same model to RNA-seq data (the DTs then represent sequencing-dependent biases). From the RNA model we predicted the expected gene-specific RNA abundance and used it as an offset term.

In all figures showing DTs, point estimates as returned by *glm4* and relatively expressed in log are plotted. The reproducibility of those point estimate accross biological samples is shown in Fig. S3C.

*Negative binomial (NB) noise model*. *θ_s_* was taken as a sample specific parameter and was empirically estimated from the variance-mean relationship expected for a NB. Specifically, for each gene we used pairs of adjacent codons occurring more than once, the rationale being that according to our steady-state model, the counts observed on the multiple instances behave as replicates (i.e. the counts are sampled from the same NB). For those pairs of codons, we computed the respective mean and variance of counts and since we assumed that *θ_s_* only depends on the sample, we then estimated *θ_s_* globally (from all pairs on all genes) by linear regression using Eq.1 (Fig.S1 E). The fit was performed using the *glm4()* function from the R package *MatrixModels* with the noise family *negative.binomial(θ_s_)* from the *MASS* package and with sparse design matrix option. Sequencing library size is used as an offset. RNA-Seq data is fitted (when available) and read counts are then predicted at every positions and used as an offset.

#### Differential expression in *ad libitum* fed vs. fasted mice

Two outlier samples (ZT12/FA/CHX and ZT04/FA/NOCHX) were excluded for the differential expression analysis and DT modelling. Statistics were computed using EdgeR [90] comparing a model including factors for time, feeding, and drug conditions against a model without the feeding term.

#### Animal experiments

Animal studies were approved by the local ethics committee, and all protocols were approved by the Service Vétérinaire Cantonal (Lausanne, Switzerland) under license VD3613. 8 weeks old male C57BL6/J mice (Charles River Laboratory) are kept under diurnal lighting conditions (12-h light, 12-h dark) at a temperature of 21 °C +/- 2 °C. After a complete night of fasting, the mice were kept without access to food for an additional period of up to 24 hours. During this time period animals were sacrificed every 8 hours starting at ZT4. Control animals were kept on *ad libitum* feeding regimen.

#### Ribosome profiling

Samples preparation for RP was performed as described in [51] except for the conditions without cycloheximide (CHX) in which fresh livers were directly lysed in ice-cold lysis buffer without CHX and directly flash-frozen in liquid nitrogen. To limit possible bias due to footprint size selection related to different conformations of the ribosome [69] [38], a larger band was cut on the TBE-gel. Libraries were generated using NEXTflex Small RNA Sequencing Kit v3 (bioo scientific) following the manufacturer’s protocol. Samples were pooled based on the Illumina indices used. Denaturated pools were spiked with 5% PhiX and clustered (loading equivalent to 3 samples per lane) onto a rapid single-end v2 flowcells at a concentration of 8pM. Libraries were sequenced on a HiSeq 2500 (Illumina) for 50 cycles.

#### (Aminoacyl-) tRNA profiling

The tRNA profiling protocol was adapted and modified from [9]. We tested the initial protocol [9] on mouse liver samples but the results showed a high proportion of unspecific ligations between the left and right probes from distinct tRNAs. We solved this issue by inverting the order of two steps in the protocol: we performed the pull-down and cleaning on magnetic beads before the splint ligation between the two DNA probes on the tRNA (Fig. S7A). Oxidation of 3’-tRNA by periodate was adapted from [60]. All the steps were performed under cold and acidic conditions to avoid deacylation of the tRNAs before Na periodate oxidation.

#### Probe Design

DNA probes were designed to target all the annotated mouse tRNAs from http://gtrnadb.ucsc.edu/. The database contains tRNA gene predictions by tRNAscan-SE [61]. tRNA sequences for *Mus musculus* (GRCm38/mm10) were downloaded and spliced *in silico*. The sequences were split in the middle of the anticodon in order to design left and right probes. After reverse complementation of the sequences, overhangs (for PCR primer binding) and unique molecular identifiers (UMIs, 2×6N) were added (right-probe adapter:

5’-GCACCCGAGAATTCCANNNNNNTGG-3, left-probe adapter:

5’-NNNNNNGATCGTCGGACTGTAGAACTC-3’). Left probes were ordered with a 5’-phosphate to allow ligation with the right probe upon annealing with the corresponding tRNA. The random nucleotides were ordered as «high fidelity wobble »to ensure homogeneous representation of the four bases in the UMI and to avoid bias. DNA probes were ordered at MicroSynth AG (Switzerland).

#### tRNA extraction and oxidation

50-100 mg of frozen mouse liver tissues were weighted under cold conditions. Beating beads were added and the samples were homogenized in 350 *µ*l of cold Qiazol (Qiagen) lysis reagent in a TissueLyser (Qiagen) for 2 x 2 min at 20 Hz. Tubes were left 5 min at room temperature. 140 *µ*l of *CHCl*_3_ was added and homogenates were shaken vigorously followed by centrifugation at 4°C for 15 min (12’000 x g). The upper aqueous phase was carefully removed and 1 volume (350 *µ*l) of buffered phenol (Phenol:chloroform:isoamyl alcool, 25:24:1, pH 4.9) was added. Samples were mixed and centrifuged for 15 minutes at 4°C (12’000 x g). Upper phase (300 *µ*l) was supplemented with 1 volume (300 *µ*l) of cold isopropanol, precipitated 30 minutes at 4°C and then centrifuged for 15 minutes at 4°C (12’000 x g). RNA pellets were dried at room temperature and re-suspended in 500 *µ*l of Sodium Acetate buffer pH 4.9 (0.2M). Samples were split in two tubes (2 x 250*µ*l) for sodium periodate oxidation (NaIO_4_) or control (NaCl) treatment. 50 *µ*l of NaCl (0.3M) or NaIO_4_ (0.3M) was added and samples were incubated for 30 minutes at room temperature. The reaction was then supplemented with 300 *µ*l Ethanol (70%) and loaded on a miRNeasy column (Qiagen). tRNA were extracted following the miRNA easy protocol from Qiagen. 390 *µ*l Ethanol (100%) was added to the flow through and loaded on a MinElute column (Qiagen). Columns were washed following the manufacturer’s protocol and RNAs were eluted in 15 *µ*l RNase-free *H*_2_O.

#### Deacylation

Purified tRNAs (14 *µ*l) supplemented with 6 *µ*l of Tris-HCl (pH 8) were deacylated by heating at 40°C for 35 minutes. Reaction was stopped by the addition of 30 *µ*l NaAcetate (0.3 M). RNAs were purified using RNA Clean & Concentrator -5 kit (Zymo) according to manufacturer’s instructions and eluted in 15 *µ*l RNase-free *H*_2_*O*.

#### 3’-tRNAs biotinylation

3’-tRNAs biotinylation was adapted from Pierce RNA 3’-End Biotinylation Kit (Thermo Fisher). Deacylated tRNAs were denaturated in 25% DMSO at 85°C for 5 minutes and directly chilled on ice. Biotinylation was performed in a 90 *µ*l reaction with 6 U of T4 ssRNA Ligase (NEB), 4 *µ*l Biotinylated Cytidine (Thermo Fisher, 1mM), 2 U RNase inhibitor, 9 *µ*l RNase Buffer (NEB), 9 *µ*l ATP (NEB, 10mM), 40 *µ*l PEG 800 (50%), and 20 *µ*l denaturated RNAs. The reaction was performed overnight at 16°C. Biotinylated tRNAs were cleaned using RNA Clean & Concentrator -5 kit (Zymo) according to manufacturer’s instructions and eluted in 20 *µ*l *H*_2_0.

#### Probes hybridization

DNA probes were synthesized by *Microsynth AG* and resuspended at a 100 *µ*M concentration. The 606 probes were then mixed at an equimolar ratio (0.15 *µ*M each) and aliquoted for further usage. Hybridization of probes was performed in a 300 *µ*l*−*reaction with 45 *µ*l probes mastermix, 30 *µ*l hybridization buffer 5x (500*µ*M EGTA, 25mM NaCl, 50mM Tris*−*HCl), 205 *µ*l RNase-free water and 20 *µ*l tRNAs. After a 15 minutes denaturation at 95 °C, the mixture was slowly cooled down to 55 °C (0.2 °C/second) and incubated for 30 minutes.

#### Beads purification

200 *µ*l of Dynabeads MyOne Streptavidin C1 (Thermo Fisher) were washed following manufacturer’s instructions for RNA usage. 250 *µ*l of beads, re-suspended in washing buffer 2x (10 mM Tris*−*HCl, 1 mM EDTA, 2M NaCl), were incubated with 300*ul* of the resulting RNA-DNA hybridization reaction for 40 minutes with gentle rotation. Beads were washed/magnetized three times with 1ml of washing buffer (1x) and re-suspended in 300 *µ*l *H*_2_*O*.

#### RNA-DNA hybrid ligation

Bead purified DNA-RNA hybrid on beads were ligated at the anticodon nick by a combination of SplintR and T4 DNA ligases (NEB) to minimize ligation efficiency bias. 300 *µ*l of DNA-RNA hybrids were splint-ligated with 2.5 U of SplintR DNA ligase and 30*µ*l of SplintR DNA ligase buffer (10X, NEB) for 1 hours at 25 °C. Then, 10 U of T4 DNA ligase (NEB) and 33*µ*l of T4 DNA ligase buffer (10x, NEB) were added. Ligation was performed overnight at 16 °C.

#### RNA digestion

Beads were magnetized and washed once with washing buffer (1X) to remove any remaining ligases. Next, beads were re-suspended in 10*µ*l *H*_2_*O*. 2 U of RNase A (Thermo Fisher) and 10 U of RNase H (NEB) with RNase H buffer (10X) (NEB) were added and digestion was performed for 30 minutes at 37 °C. Elution buffer (5X) was added for a final concentration of 50 mM tris pH8, 10 mM EDTA, 1% SDS and samples were incubated at 65 °C for 30 minutes with intermittent shaking to retrieve ligated DNA probes. Beads were magnetized and supernatant extracted. DNA ligated probes were purified using DNA Clean & Concentrator -5 kit (Zymo) according to manufacturer’s instructions and eluted in 20 *µ*l RNase-free *H*_2_*O*.

#### qPCR for quality control and relative concentration estimation

The relative concentration of the resulting DNA ligated-probes was assessed by quantitative PCR (qPCR) using the LightCycler 480 SYBR Green I Master kit (Roche). 3.5 *µ*l of 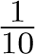 diluted samples was used to assemble a 10 *µ*l reaction and run on a Light Cycler 480 II (Roche) with the primers CAGAGTTCTACAGTCCGACGAT and TTGGCACCCGAGAATTCCA (matching each probe’s ends) at a final concentration of 0.3 *µ*M. Cycling conditions consisted of an initial denaturation step of 5 min at 95 °C followed by 40 cycles of 30 s at 95 °C and 45 sat 45 °C. The Cp obtained were used to calculate the optimal number of PCR cycles amplification required for library amplification, as described previously [51]. The number of required cycles were between 13 and 17 depending on the experiments and samples. The quality of the ligation was assessed on a Bioanalyzer small RNA chip (Agilent Technologies).

#### PCR amplification

The PCR was designed following Illumina’s recommendation taking advantage of the indexed oligos from the TruSeq small RNA kit. A 50 *µ*l-reaction was assembled with Kapa Polymerase and 15 *ul* of DNA ligated probes, and run for the optimal number of PCR cycles calculated as described above.

#### Library postprocessing

Amplified libraries were purified with 100 *µ*l AMPure XP beads (Beckman) and eluted in 20 *µ*l resuspension buffer (10 mM Tris, pH8.0). Libraries were quantified with Picogreen (Life Technology) and usually yield 50-400 ng DNA. The libraries size patterns were checked using a fragment analyzer (Agilent).

#### Library Sequencing

An equimolar library pool was prepared from the individual libraries based on the concentrations calculated from Picogreen data. Pools were denaturated with NaOH and neutralized with HT1 buffer (Illumina) to reach a final concentration of 20pM. Pools were spiked with 10 % PhiX and clustered (loading equivalent to 12 samples per lane) onto a rapid single-end flow cell v2 at a final concentration of 7pM. Sequencing was performed on a HiSeq 2500 (Illumina) in rapid mode for 130 cycles.

#### Data Analysis

To assess the fidelity of left/right probe ligation and efficiency of hybridization, a fasta file with all the possible combinations between left and right probes was created. In case two tRNA genes share the same left/right probes, they were grouped and annotated accordingly. It led us to a total of 68526 sequences. Genome index was generated with STAR with options

`–runMode genomeGenerate –genomeSAindexNbases 3`. Fastq files were trimmed for adapters. Sequencing reads were aligned against the index with parameters:

`–outFilterScoreMinOverLread 0 –outFilterMatchNminOverLread 0`

`–winAnchorMultimapNmax 1000 –outFilterMismatchNmax`

`–clip3pNbases 6 –clip5pNbases 6`

`–outFilterMultimapNmax 50 –outSAMattributes NM nM NH NM –alignIntronMax 1`

`–alignEndsType EndToEnd –seedSearchStartLmax 20 –seedMultimapNmax 100000.`

For each combination, the number of counts were computed and corrected for PCR duplicates using both unique molecular identifiers sequences. Reads larger than 60 nucleotides with less than 3 mismatches and less than 5 insertions compared to the reference were retained. Combinations with less than 10 reads were discarded. Reads mapping to combinations of probes coming from different codons were reassigned in function of the newly ligated sequence. Abundances of tRNA coding for the same codon were summed up and normalized by library size using *edgeR* R package [90]. Because tRNA moieties (the two halves of the tRNA) have very similar sequences, and since the specificity of the hybrid DNA-RNA around the anticodon is important for the ligation, we used the sequence around the anticodon to reassign the ambiguous combinations.

### Data Availability and specificity

Sequencing data of this study have been submitted to Gene Expression Omnibus (GEO) under accession number GSE126384: https://www.ncbi.nlm.nih.gov/geo/query/acc.cgi?acc=GSE126384). Datasets and GEO references: Atger Liver (GSE73553, n=84), Howard Liver (SRR826795, SRR826796, SRR826797, n=3), Huh Linter (SRR5227294, SRR5227295, SRR5227296, SRR5227303, SRR5227304, SRR5227305, n=6), Jan Yeast (SRR1562907, SRR1562909, SRR1562911, SRR1562913, n = 4), Kidney Castelo (GSE81283, n=24), Guydosh Yeast (SRR1042865, SRR1042866, SRR1042867, n= 1).

## Supporting information

Single-codon dwell times

Codon-pair dwell times

Ribosome profiling in AL and FA mouse liver

(aminoacyl-) tRNA profiling in AL and FA mouse liver

## Acknowledgments

We thank Nicolas Bonhoure and Stefan Morgenthaler for useful discussions. Research in the Naef laboratory was supported by the EPFL. Some computations were performed on the Vital-It computing platform.

## Supplementary material

Table 1. Inferred single-codon DTs

Table 2. Inferred codon-pair DTs

Table 3. Ribosome profiling in AL and FA mice w/o CHX

Table 4. (aminoacyl-) tRNA profiling in AL and FA mice

## Code availability

Snakefile to model DTs and flux from ribosome profiling data will be made publicly available on github upon publication.

---

**Figure S1:**
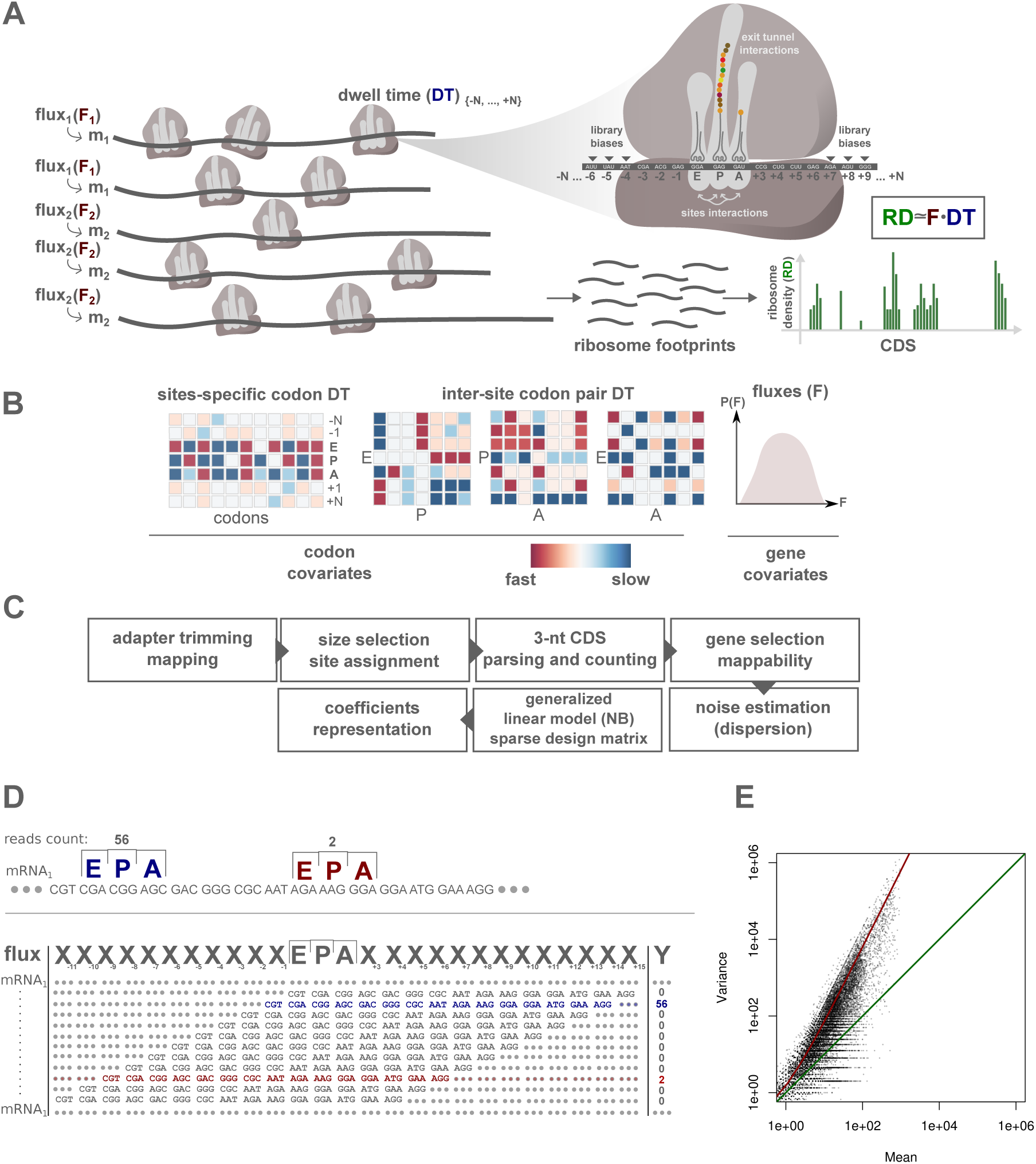
Modeling ribosome fluxes and codon-specific DT including codon-pair contributions. A. Ribosome elongation: Snapshot of two different mRNA species (*m*_1_*, m*_2_) translated with two different fluxes (*F*_1_*, F*_2_). Zoomed ribosome shows that numerous factors regulate ribosome DTs: 40 codons (+20,-20) around the E site are taken into account in the model to alleviate possible library biases, exit tunnel interactions, and influence of upstream/downstream sequences. Codon-pair interactions between the three sites (E, P, A) are also modeled. The ribosome densities on the mRNAs are estimated by ribosome profiling, and modeled as genes fluxes multiplied by DTs. B. Single-codon and codon-pair DTs are visualized in a heatmap, relative to the position mean. The matrix of codon-pair interactions (E:P, P:A, and E:A) shows all possible combinations of codon pairs. All regression coefficients (gene fluxes and DTs) are inferred genome-wide. C. Bioinformatics pipeline: Sequencing reads are trimmed and mapped to the genome. Reads are first selected based on their size and E sites are assigned for each read. All annotated CDSs are parsed with a step of 3 nucleotides and number of reads are reported at each position. Genes with insufficient total read counts and read densities are removed, as well as regions with non-unique mappability. Dispersion parameters for negative binomial distributions are estimated for each sample and the GLM is fitted with a sparse design matrix and negative binomial (NB) noise model. DTs are centered (in log2 scale) and represented as shown in Fig. S1B. D. Construction of the data matrix for the GLM. Example of a gene CDS with two different positions (dark blue and dark red) covered by 56 and 2 reads, respectively. The assigned E, P, and A sites are shown. The CDS is parsed 3-by-3 and a matrix is designed with the corresponding position-dependent codons. E. Mean and variance of measured counts for pairs of codons occurring multiple times on a gene. The green line shows a *Poisson* regime with the variance equal to the mean. The red line represents the estimated fit for a negative binomial distribution (Methods). The dispersion parameter is estimated from these fits and used to parameterize the NB used in the GLM, independently for each sample.

**Figure S2:**
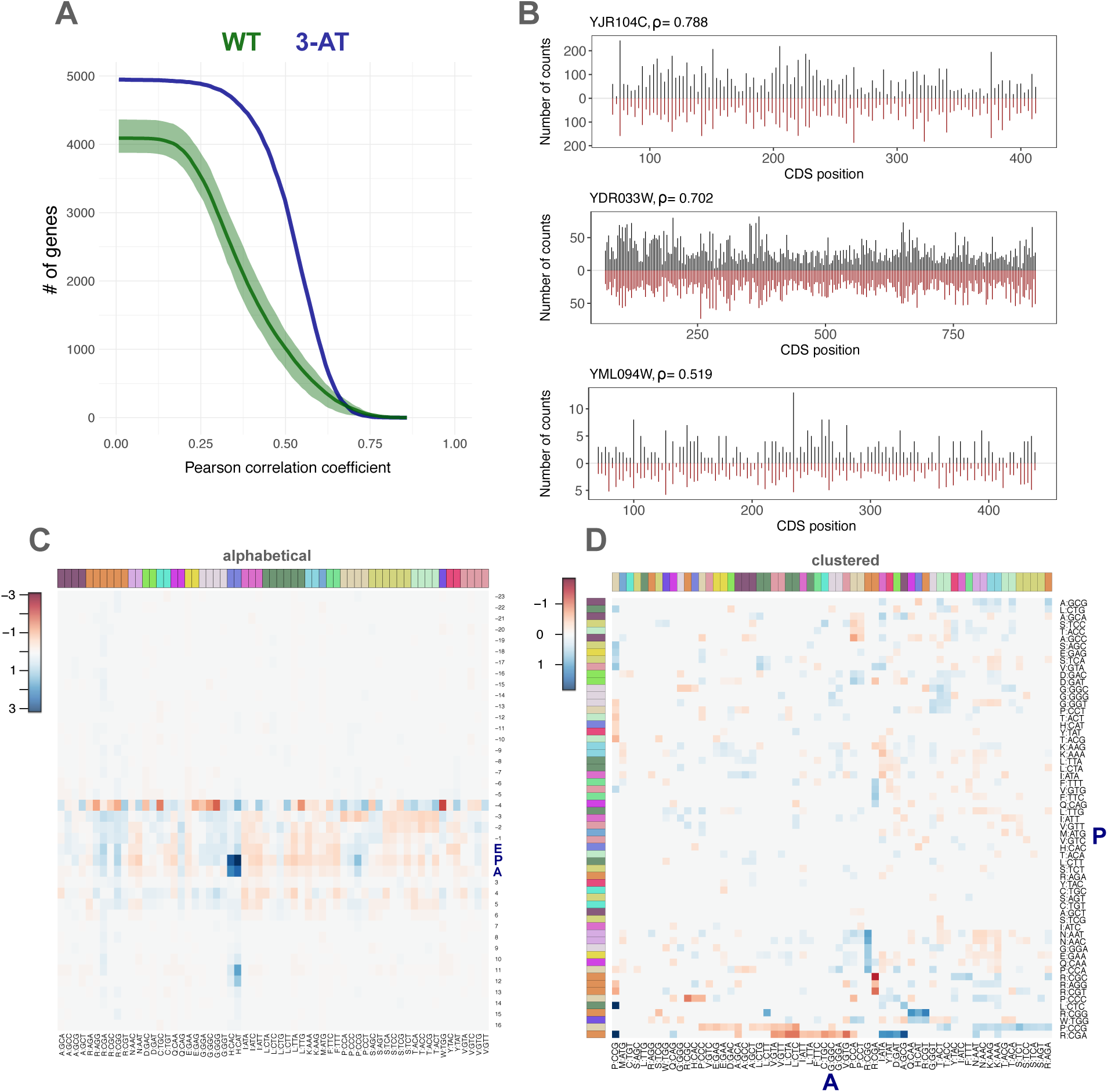
In yeast ribosome DT anticorrelate with codon usage and exhibit codon-pair interactions. Panels for the single-codon DTs are retrieved from the fit with the P:A interaction. A. Gene-specific Pearson correlation coefficient (*ρ*) between measured and fitted read counts. Number of genes with *ρ* larger than x-axis values are depicted. WT (green) and 3-AT (blue). B. Measured (black) and fitted (red) read counts for three representative genes. C. Heatmap representation of the DTs (log2, mean centered per site) in a window of 40 codons around the E site. Codons are ordered by amino acid and colored accordingly at the top of the heatmap. DTs with p *>*= 0.05 are not shown(set to zero). D. Interaction matrix for the pairs P:A (log2). Codons are colored according to amino acid. Codons in both sites are hierarchically clustered based on the euclidean distance matrix and a complete linkage algorithm. Fast and slow interactions are shown respectively in dark red and dark blue (colorbar). DTs with p-value *>*= 0.05 are set to zero.

**Figure S3:**
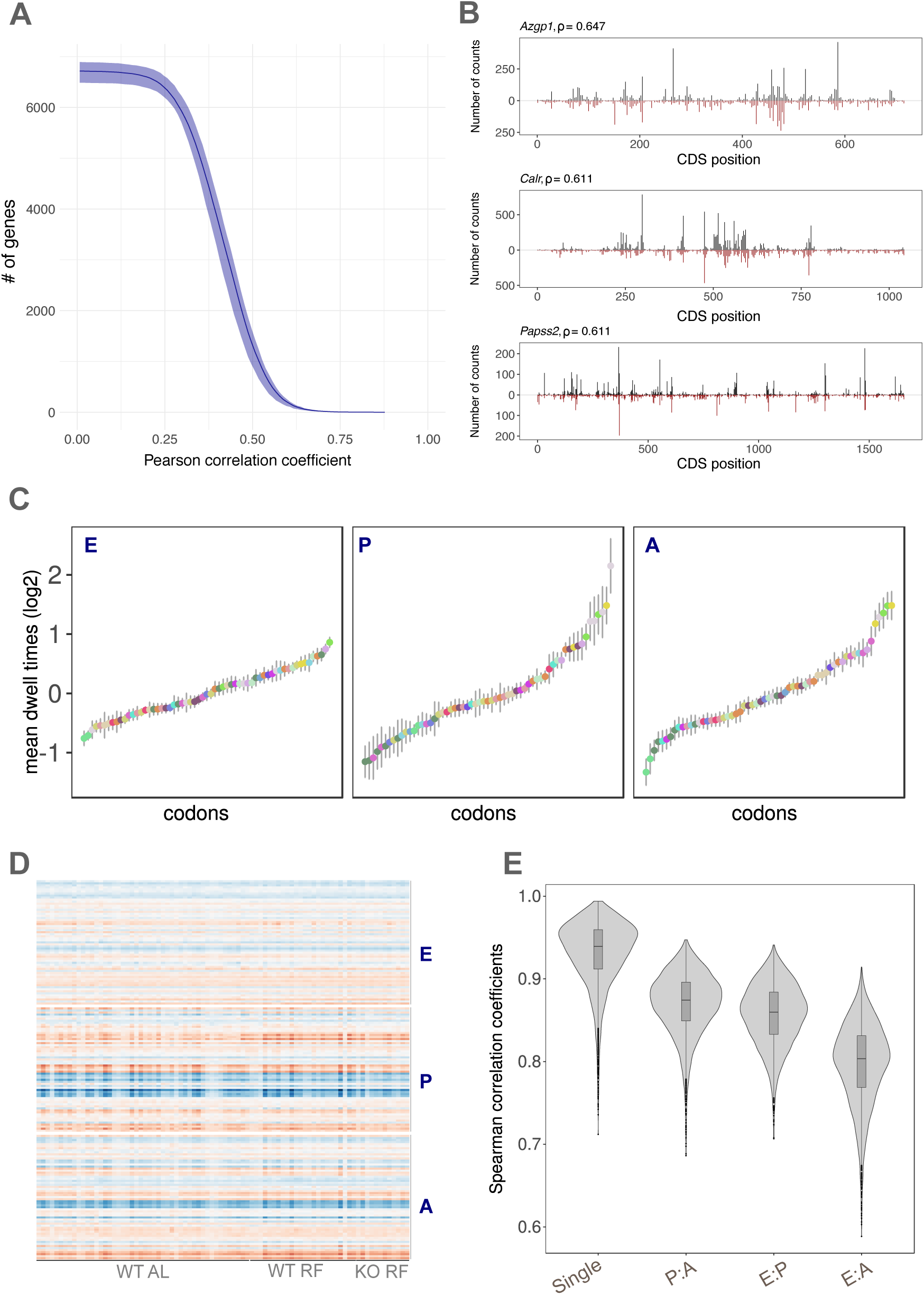
Sample specific DTs and correlations. A. Gene-specific Pearson correlation coefficient (*ρ*) between measured and fitted read counts for four AL samples [51]. Number of genes with *ρ* larger than x-axis values are depicted. B. Measured (black) and fitted (red) read counts for three representative genes in one AL sample. C. Mean DTs and standard deviation over the 84 samples for the E, P, and A sites. D. DTs (log2, mean centered per site) at the E, P, and A sites for the 84 samples in the three conditions WT AL (WT ad libitum), WT RF (WT night-restricted feeding), and KO RF (*Bmal1* KO night-restricted feeding). E. Inter-sample Spearman correlation coefficients for site-specific DTs (single) and codon-pair DTs for the interactions P:A, E:P, and E:A.

**Figure S4:**
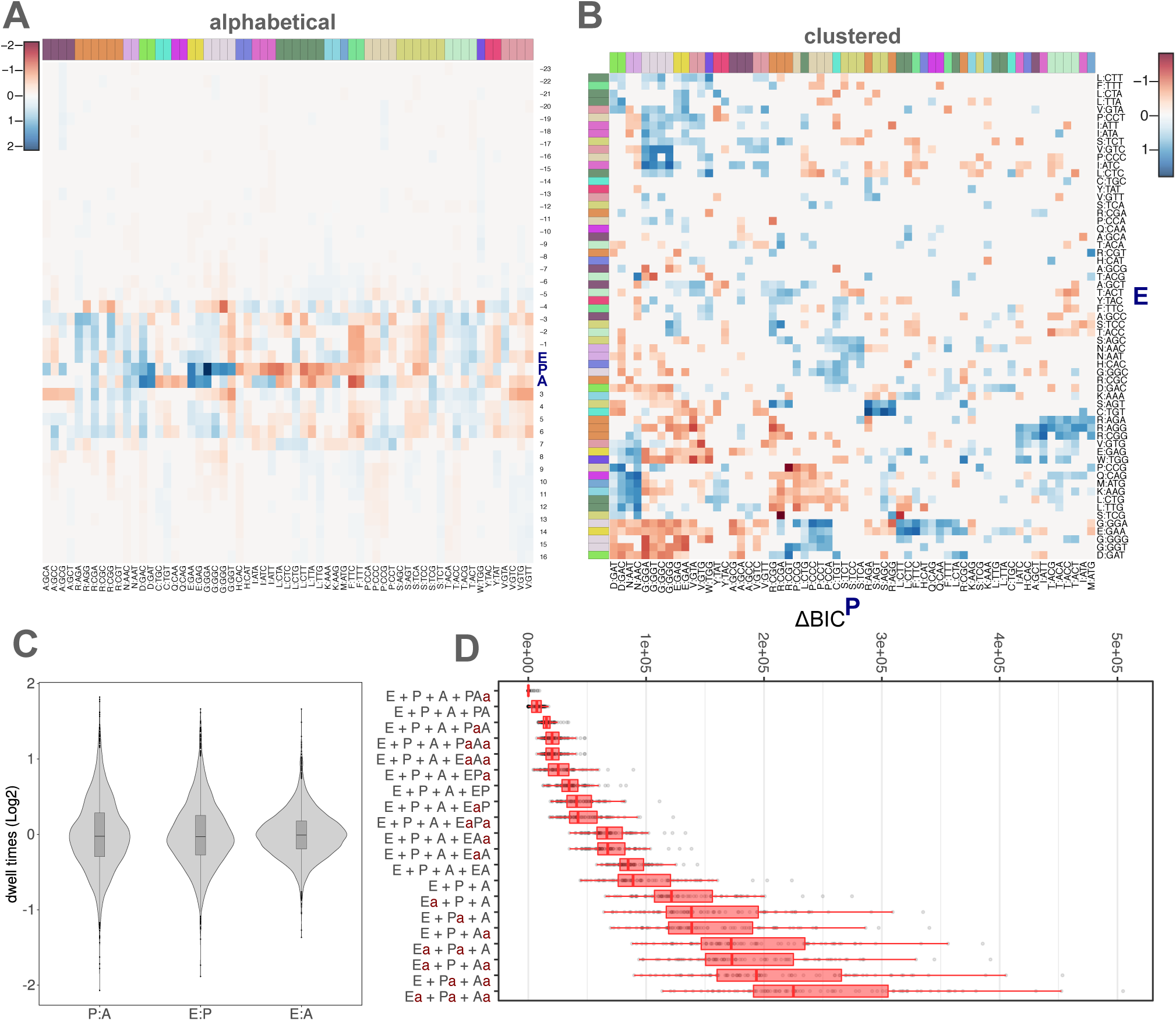
Extended analysis of the translation elongation landscape in mouse liver. Single-codon DTs are retrieved from the fit with the P:A interaction. A. Heatmap representation of the DTs (log2, mean centered per site) in a window of 40 codons around the E site. Codons are ordered by amino acid and colored accordingly. DTs with *p >*= 0.05 are not shown (set to zero). B. Interaction matrix for E:P interactions (log2). Codons are colored according to amino acid. Codons in both sites are hierarchically clustered based on the euclidean distance matrix and a complete linkage algorithm. Relatively fast and slow interactions are shown respectively in dark red and dark blue. DTs with p-value *>*= 0.05 are not shown (set to zero). C. DT (log2) distributions and boxplots for the three interaction terms P:A, E:P and E:A of the 84 samples in mouse liver. D. Differences of Bayesian Information Criterion (ΔBIC) between the model shown and the best model. (ΔBIC) is computed for each sample and proposed model, in which the alphabet for the DT covariates was taken as either the 20 natural amino acids or the 61 sense codons. A lowercase ‘a’ on the right of an uppercase letter indicates that the amino acid alphabet was used for this position.

**Figure S5:**
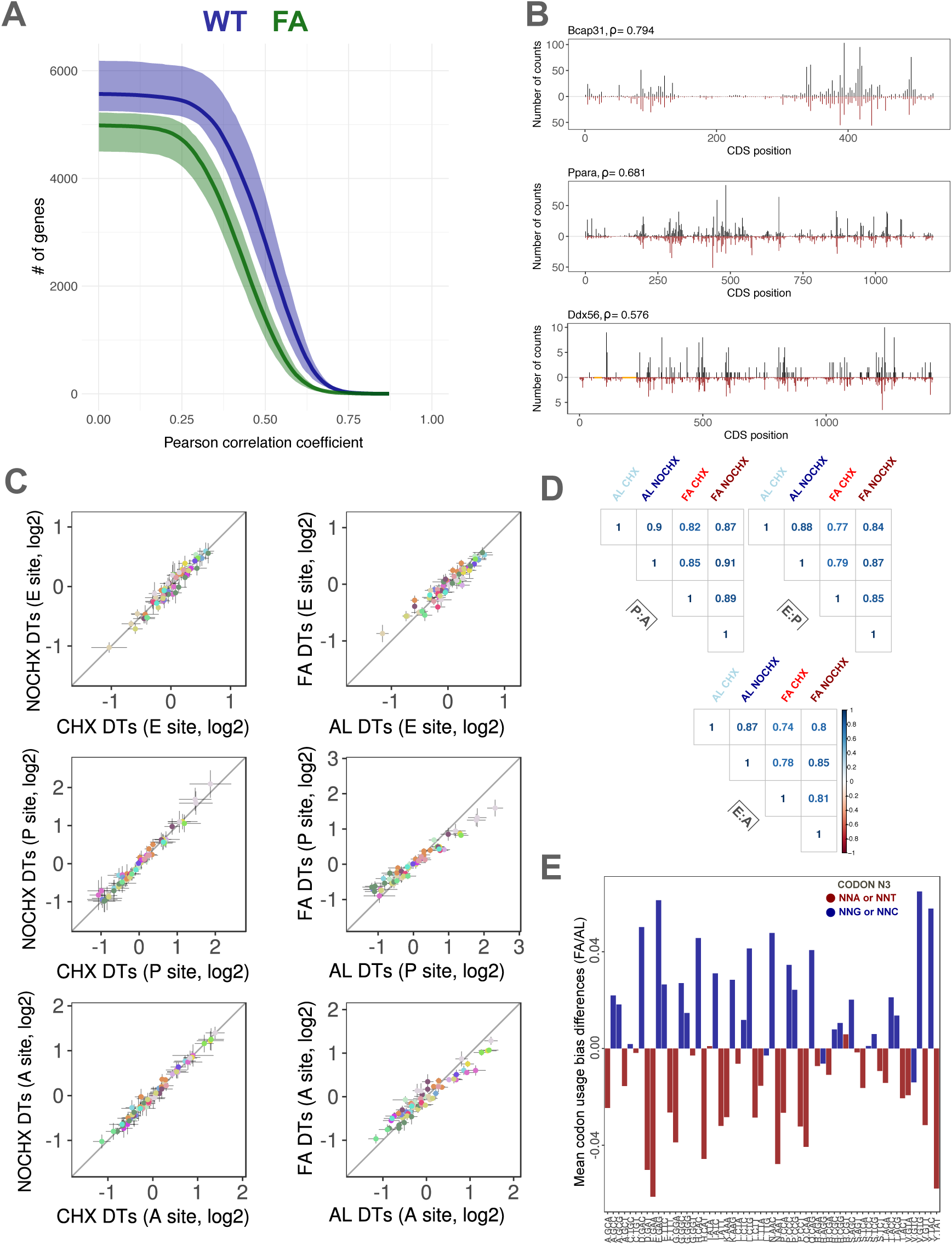
Ribosome DTs are not affected in fasted mice and with cycloheximide. A. Gene-specific Pearson correlation coefficient (*ρ*) between measured and fitted read counts. Number of genes with *ρ* larger than x-axis values are depicted. WT (blue) and FA (green) B. Measured (black) and fitted (red) read counts for three representative genes in one AL sample. C. DTs in the E (top), P (middle) and A site (bottom) for the CHX vs. NOCHX conditions (left) and AL vs. FA conditions (right). Standard deviation is depicted. D. Pearson correlation coefficient between the different conditions for the interaction terms (log2) E:P, P:A, and E:A. E. Codon usage bias is computed for each gene up- or down-regulated in fasted animals and averaged. The difference in codon usage bias is computed between the FA and AL conditions. Codons are colored accordingly to their nucleotide at the third position (G-C in blue and A-T in red).

**Figure S6:**
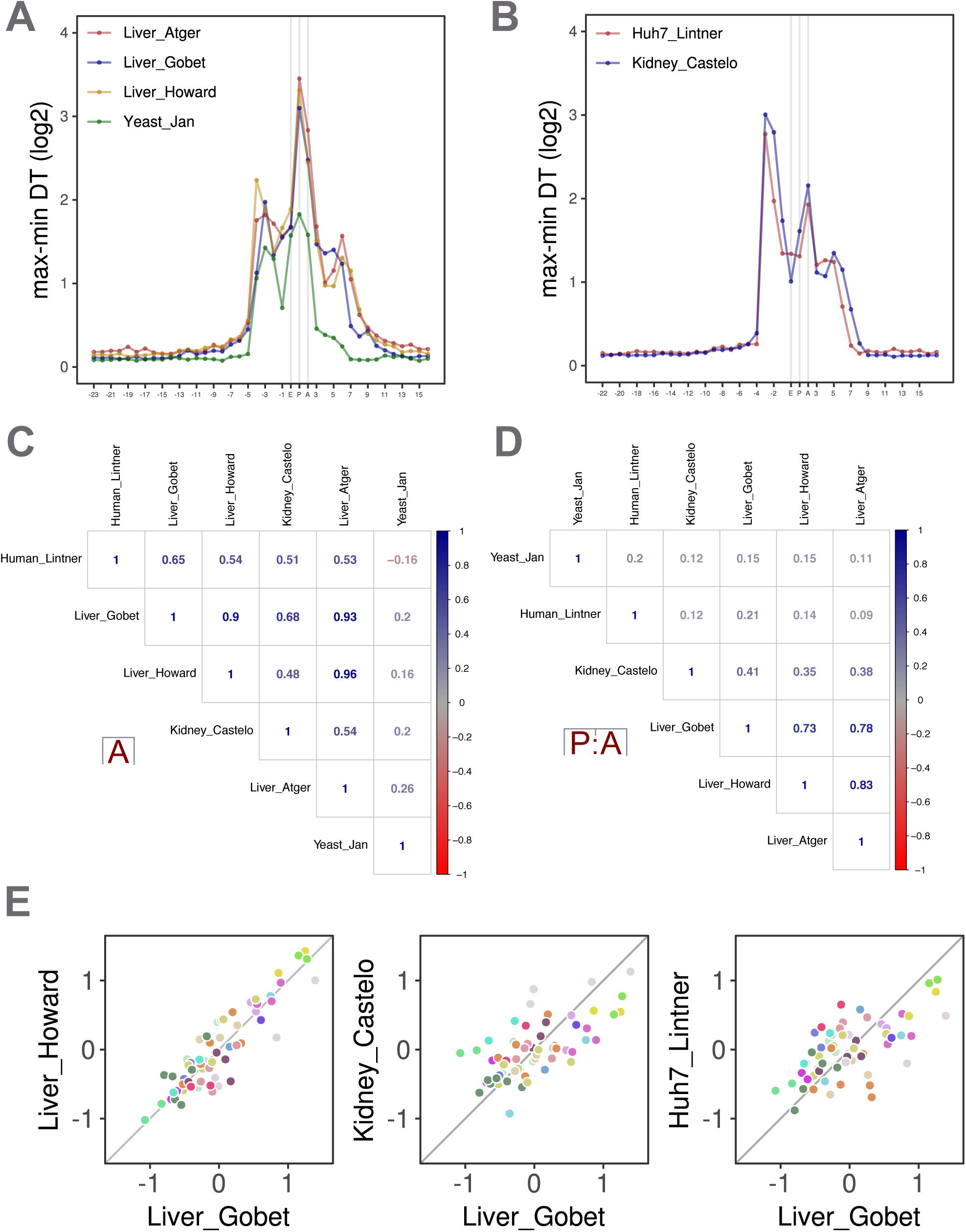
Meta-analysis revealed conserved DT patterns and technical biases. Analysis of published ribosome profiling datasets in yeast (Yeast Jan) [49], mouse liver (Liver Atger, Liver Howard, Liver Gobet (this paper)) [51, 57], mouse kidney (Kidney Castelo) [58] and in a human hepatocyte cell line (Huh7 Lintner) [59]. DTs were inferred for each sample and averaged by condition. A. Spread of the DTs (max-min, log2) at every positions in a window of 40 codons around the E site for studies using small RNA library protocols. Colors show the different datasets. B. Same as (A) for studies using “cDNA circularization” library protocols. C. Correlation for the A site DTs between the different datasets (Pearson coefficient is color coded). D. Same as (C) for P:A interaction. E. DTs in the A site for the Liver Howard, Kidney Castelo and Huh7 Linter datasets vs. DTs in the A site from the RP data in this paper (Liver Gobet, AL and FA).

**Figure S7:**
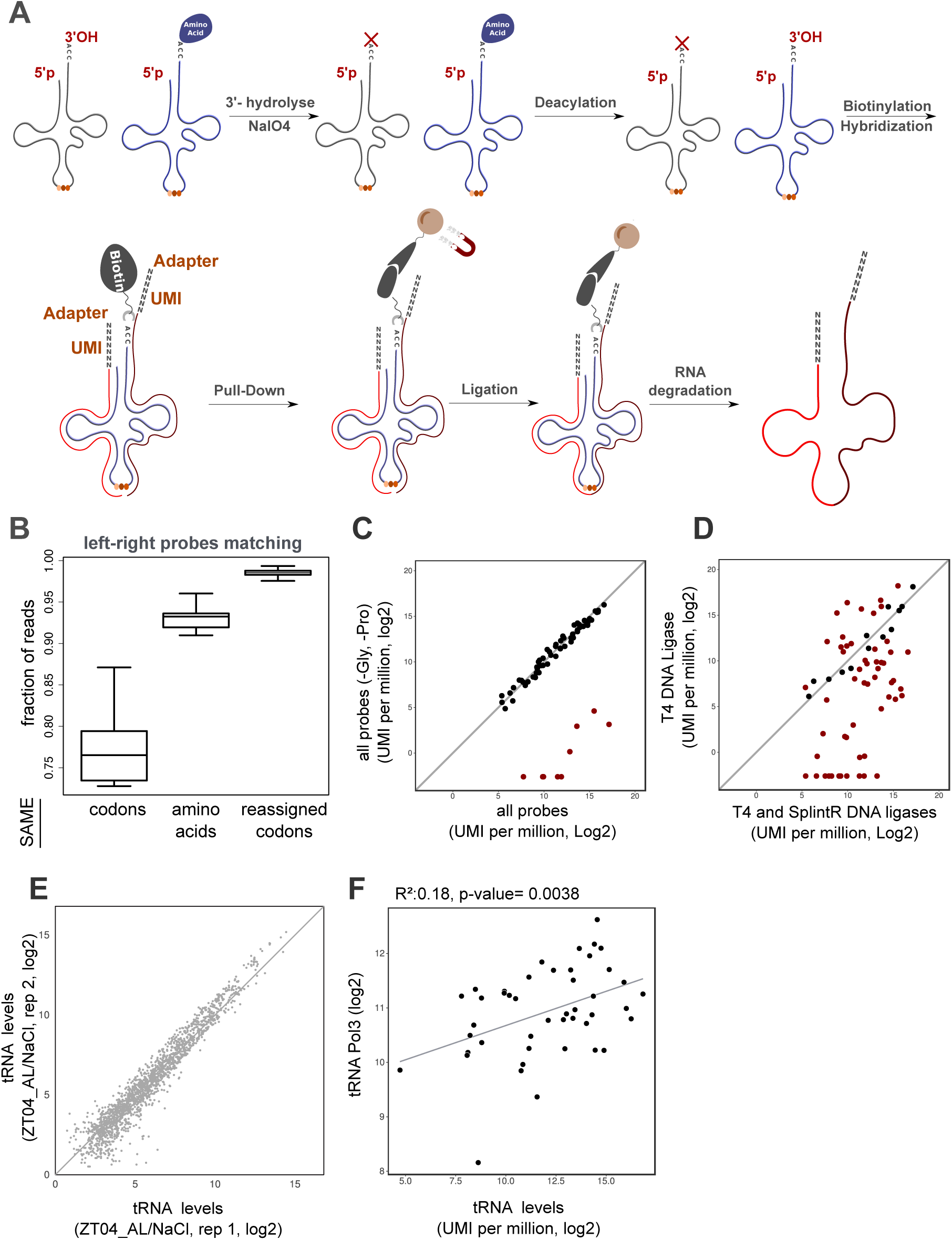
(aminoacyl-) tRNA profiling in AL and FA mice (bis) A. tRNA profiling protocol. tRNAs were extracted in acidic conditions and uncharged tRNAs were hydrolyzed at 3’-end upon periodate treatment. tRNAs were treated with NaCl in control conditions. Then, tRNAs were deacylated and biotinylated at their 3’-end. Mix of left and right DNA probes were hybridized to the tRNA pools and pulled-down on magnetic beads through biotin-streptavidin interactions. Nicks in the anticodon between the left and right probes were ligated. tRNAs were degraded and DNA probes sequenced after amplification. B. Reads were mapped on every combination of left and right probes. Fraction of reads corresponding to left-right probe combinations belonging to the same codon or amino acid is reported for the 24 samples. The same measure is computed after reassignment of the probe combinations (Methods). C. tRNA abundances (log2) at the codon level for the control vs. altered conditions in which probes related to tRNAs coding for Pro and Gly were removed. D. tRNA abundances (log2) at the codon level for experiments with T4 or SplintR DNA ligases. Significant differences are shown in red. E. tRNA abundances (log2) at the probe level between biological replicates in the NaCl/AL condition at ZT04. F. Correlation between tRNA abundances in control AL vs. RNA polymerase III (POL3) ChIP-Seq signal quantified on the tRNAs gene loci. Data were extracted from the supplementary table of ref. [62].

**Figure S8:**
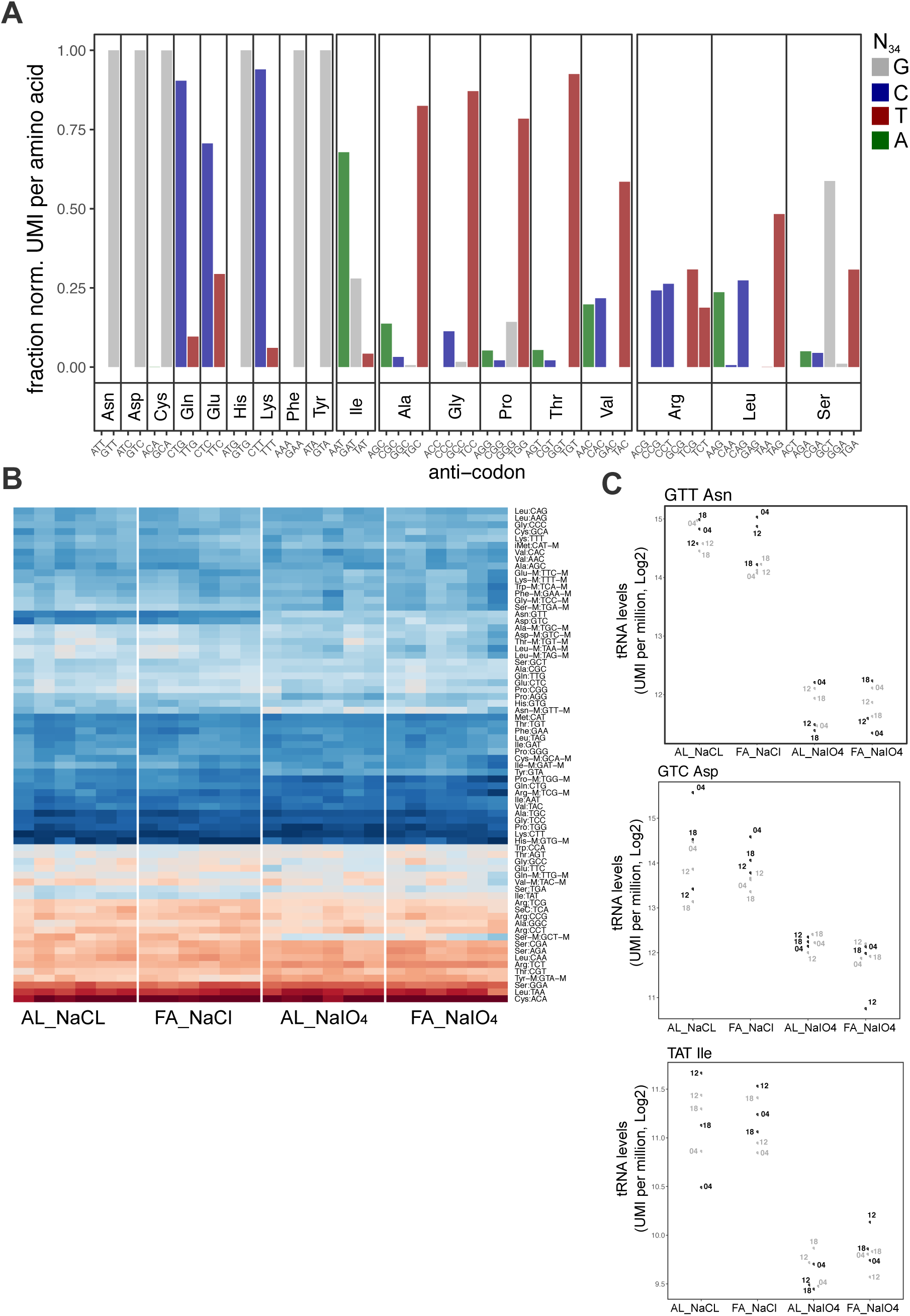
(aminoacyl-) tRNA profiling in AL and FA mice (bis) A. Fraction of reads (UMI) per amino acid for the different isoacceptor tRNAs. Colors indicate the nucleotide in position 34 on the tRNA. B. Heatmaps of the normalized tRNA read counts (log2) at the codon level for the 24 samples. tRNAs are ordered with hierarchical clustering. Dark blue (high), dark red (low). C. Expression levels for three tRNA coding respectively for Asn, Asp, and Ile, showing 5-8 fold differences between control and periodate-treated conditions.

